# FLASH X-rays preserve neutrophil chromatin to boost antitumor immunity

**DOI:** 10.64898/2026.10.11.758276

**Authors:** Xiaolin Shi, Si Qiu, Yiwei Yang, Yongchao Jiang, Jianxin Wang, Xiaofeng Ding, Yufan He, Meiqi Lv, Hongyan Li, Qing Li, Baosheng Li, Zheng Zhou, Kui Zhou, Dexin Xiao, Peng Li, Chuanyu Cao, Lijun Shan, Yu Liu, Tingting Wang, Feng Gao, Caiyong Ye, Congchong Yan, Ji Huang, Yuan Wang, Guangsheng Shi, Fengjuan Jiang, Yuqi Xiao, Hongdao Qu, Xun Jin, Jianghong Zhang, Yan Pan, Yang Bai, Feng Xi, Liang Wu, Xiaobo Du, Jianheng Zhao, Jian Wang, Min Xie, Dai Wu, Chunlin Shao, Haowen Zhang

## Abstract

Ultrahigh dose-rate FLASH radiotherapy spares normal tissue while preserving antitumor activity, positioning it as a promising technology in radiation oncology, but its immunomodulatory mechanisms remain elusive. Here, using a bilateral subcutaneous tumor model in mice, we show that 8-Gy FLASH X-rays, versus an equal dose of conventional dose-rate (CONV) X-rays, elicit a robust abscopal antitumor effect with anti-PD-1 therapy. Single-cell transcriptome and T cell receptor (TCR) profiling reveal that FLASH X-rays plus anti-PD-1 enhance antitumor immune infiltration and TCR clonal expansion in both irradiated and abscopal tumor lesions. Mechanistically, the ultrashort temporal continuity of FLASH dose delivery conserves chromatin compaction in neutrophils during exposure and inhibits neutrophil extracellular trap (NET) formation, thereby dismantling NET-mediated CD8^+^ T-cell exclusion within primary tumors. Conversely, CONV or split-dose FLASH X-rays induce marked chromatin modifications prior to dose completion, leading to chromatin hyper-relaxation which promotes NETosis and consequently abrogates systemic antitumor immunity. Notably, this detrimental effect can be reversed by pharmacologically suppressing PARylation-mediated chromatin modification and decondensation in neutrophils. These findings identify radiation-induced aberrant chromatin relaxation in neutrophils as a critical determinant of radioimmunotherapy outcomes, providing a rationale for redefining the FLASH effect and advancing its clinical translation.

## Introduction

The advent of immunotherapy has revived clinical interest in the abscopal effect, whereby localized radiotherapy can trigger regression of distant metastases. In this context, combining immune checkpoint inhibitors (ICIs) with radiotherapy has emerged as a compelling strategy, demonstrating robust efficacy against advanced metastases^1–3^ and reshaping radiotherapy practice^4,5^. However, the clinical translation remains inconsistent, attributable not only to direct radiotoxicity to intratumoral immune cells^6,7^, but also to radiation-driven immunosuppressive microenvironmental remodeling, encompassing upregulation of immunosuppressive factors such as plasminogen activator inhibitor-1 (PAI-1)^8^ and amphiregulin^9^, as well as activation of cancer-associated fibroblasts (CAFs)^10^, and induction of NETosis^11^, the latter two of which form physical barriers that restrict antitumor immune infiltration. Given these obstacles, a transformative radiotherapy modality that overcomes radiation-induced tumor immune microenvironment (TIME) dysfunction and potentiates ICI efficacy is urgently needed.

FLASH radiotherapy, delivered at an ultrahigh dose rate exceeding 40 Gy/s, represents a major conceptual and technological advance in radiation oncology, as it substantially spares normal tissues without compromising tumor control, a phenomenon termed the FLASH effect^12–17^. Its clinical promise is supported by ongoing trials (FAST-02^18^, LANCE^19^, IMPulse^20^, Flash-Skin I^21^) and the completed first-in-human study, FAST-01^22^. Beyond the sparing effect, emerging evidence suggests that, compared with conventional dose-rate (CONV) irradiation (IR), FLASH radiotherapy remodels the TIME toward a pro-immune phenotype and enhances immunotherapy efficacy^23,24^. Although our previously proposed “DNA integrity” hypothesis highlighted the critical role of instantaneous radiation delivery in mitigating the adverse effects of radioimmunotherapy^25^, the mechanisms underlying the systemic antitumor benefit of combining FLASH with ICIs remain largely elusive.

Here, we combined FLASH photons with anti-PD-1 antibody (αPD-1) to investigate how the temporal profile of dose delivery governs the neutrophil chromatin compaction, NETosis, and consequent TIME remodeling. Our work demonstrates that FLASH, when combined with αPD-1, achieves superior control of metastatic tumors over CONV X-rays, and identifies neutrophil chromatin modifications as a central mechanistic hub of the FLASH effect, bridging conceptual understanding and clinical translation.

## Results

### 1. FLASH outperforms CONV X-rays in enhancing abscopal response to αPD-1

To evaluate the abscopal efficacy of FLASH photon IR in combination with αPD-1, we delivered a single 8-Gy dose of CONV or FLASH X-rays to bilateral subcutaneous MC38 tumor-bearing mice using a horizontal beam with a uniform spatial dose distribution (Fig. 1a and Extended Data Figs. 1 and 2a). Both FLASH X-rays plus αPD-1 (FA) and CONV X-rays plus αPD-1 (CA) regimens effectively controlled primary tumors (Fig. 1b), but FA markedly outperformed CA in eradicating secondary tumors (Fig. 1c), yielding an over 4-fold increase in survival (Fig. 1d). Primary tumor recurrence was similar, whereas secondary tumors recurred in 10/12 of CA mice versus 4/12 of FA mice (8/12 remained tumor-free) (Fig. 1e,f). Mice receiving αPD-1 or IR alone all succumbed to tumor progression (0% survival rate) (Fig. 1d). Notably, the superior abscopal effect was not attributable to direct radiosensitivity differences, as FLASH and CONV IR showed comparable clonogenic survival and similar expression of *H2-K1*, *Cd274*, and high mobility group box 1 (*Hmgb1*) in MC38 cells (Extended Data Fig. 2b,c). To assess immune memory, tumor-free FA mice were rechallenged with MC38 or B16-F10 cells 90 days later (Fig. 1g). All MC38-rechallenged mice rejected tumors, whereas B16-F10 tumors grew progressively, comparable to naïve controls (Fig. 1h,i). Furthermore, splenocytes from tumor-free FA mice showed MC38-specific T-cell activation upon co-culture, with significantly upregulated CD69 and interferon-gama (IFN-γ) in both CD4^+^ and CD8^+^ T cells versus B16-F10 stimulation (Extended Data Fig. 2d and Fig. 1j–l). Thus, FA treatment induced robust abscopal antitumor immunity and durable, antigen-specific memory.

**Fig. 1.**
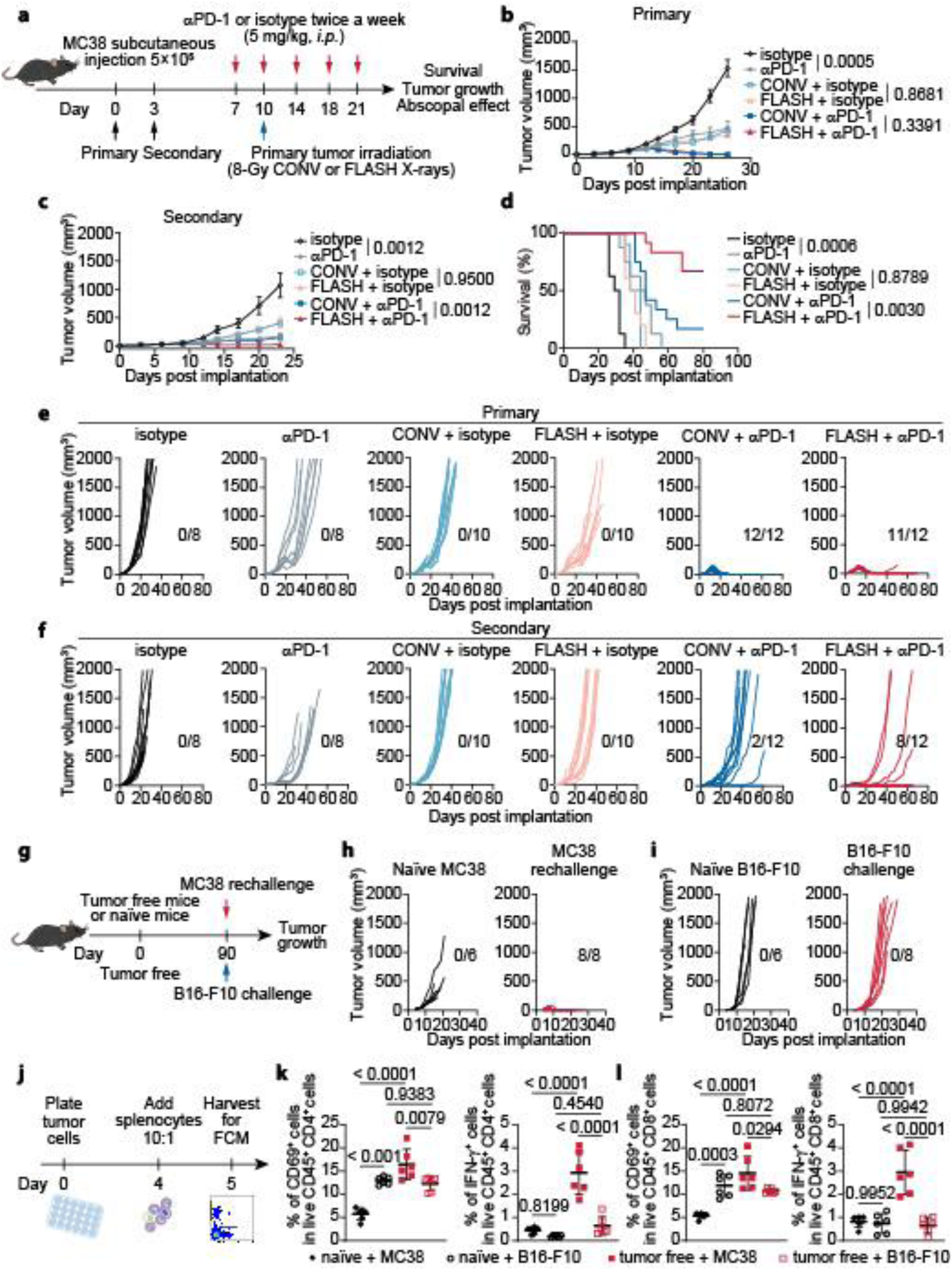
Abscopal tumor control by FLASH versus CONV X-rays with αPD-1. **a**, Treatment schema of the abscopal effect model. **b**,**c**, Tumor volumes of primary (**b**) and secondary (**c**) tumors were monitored. n = 8–12 mice per group. **d**, Kaplan–Meier survival curves of tumor-bearing mice receiving the indicated treatments was performed. n = 8–12 mice per group. **e**,**f**, Individual tumor growth curves of primary (**e**) and secondary (**f**) tumors are shown. **g**, Schema for evaluating the antitumor immune memory of tumor-free mice. **h**,**i**, Tumor growth curves of naïve (black) mice and tumor-free (red) mice (previously treated with FLASH X-rays + αPD-1) after rechallenge with MC38 (**h**) or with the unrelated B16-F10 melanoma (**i**). **j**, Splenocytes from tumor-free mice in the FLASH X-rays + αPD-1 group or from naïve mice were co-cultured with tumor cells for 24 h, then harvested for analysis of T-cell activation markers. **k**,**l**, Flow cytometry quantifications of CD69 and IFN-γ levels in CD4^+^ T cells (**k**) and CD8^+^ T cells (**l**). Ratios represent the number of tumor-free mice per treatment group and data are presented as mean ± SEM (n = 6 mice per group). Statistical significance was assessed by repeated-measures ANOVA (**b**,**c**), log-rank test (**d**), and one-way ANOVA (**k**,**l**).

### 2. FLASH X-rays bias T-cell clonal differentiation toward functional antitumor trajectories

To define the immune basis of the abscopal divergence, we profiled the transcriptomes of FA and CA tumors. Bulk RNA sequencing (RNA-seq) revealed progressive immune composition shifts, with the clearest FA versus CA divergence in distant tumors at day 6 post-IR (Fig. 2a). In these lesions, FA exhibited higher abundance of T cells, natural killer cells, and monocytes/macrophages (Extended Data Fig. 3a), elevated T-cell-recruiting chemokines (*Ccl4*, *Ccl5*, *Cxcl10*, *Cxcl11*)^26^, and inflammatory markers (*Csf2*, *Cxcl2*) (Fig. 2b). Single-cell RNA-seq of 228,350 tumor-derived cells resolved 13 major cell types and confirmed treatment-associated lymphoid and myeloid remodeling (Fig. 2c and Extended Data Fig. 3b,c). These compositional differences were not accompanied by acute DNA damage burden differences, as γ-H2AX levels at 6 h were comparable across FLASH and CONV-treated conditions in CD45^−^, CD45⁺, CD8⁺, and F4/80⁺ populations (Extended Data Fig. 3d–g). Given the central role of T cells in αPD-1-mediated antitumor immunity, we profiled T-cell clonality and transcriptional states. At day 6 post-IR, T cell receptor (TCR) sequencing showed that hyperexpanded clonotypes, undetectable in CA, were enriched in FA tumors (16.2% irradiated, 10.8% distant); large clonotypes represented 34.3% of primary and 21.2% of secondary FA tumors versus < 20% at both sites in CA (Fig. 2d). FA also displayed a higher frequency of clonotypes shared across tumor sites and expanded at one or both sites (33.0% vs 12.9% irradiated, 10.6% vs 6.3% distant) (Fig. 2e,f). In line with these sharing profiles, FTY720 blockade of lymphocyte egress^27^ attenuated the abscopal response in FA-treated mice (Extended Data Fig. 3h–j), suggesting that FA more effectively converts local IR into a trafficking-dependent systemic T-cell response. Subclustering of T cells (Extended Data Fig. 4a,b) revealed higher abundance of most subsets in FA than in CA across primary and secondary sites, whereas Migratory and interferon (IFN)-Stimulated CD8^+^ T cells showed the opposite pattern, particularly in primary tumors (Fig. 2g). Pseudotime analysis identified three trajectories sharing an early region associated with Resting/Memory and Effector-Priming states (Fig. 2h and Extended Data Fig. 4c). Path 1 progressed toward Migratory and IFN-Stimulated states dominated by IFN-response programs, whereas path 3 extended toward Activated/TRM-like and Cytotoxic states with stronger TCR signaling and effector/cytotoxic programs (Fig. 2i and Extended Data Fig. 4d–f). When projected onto these trajectories, CA cells preferentially localized to path 1, whereas FA cells were enriched in later regions of path 3, with path 2 showing limited group-specific bias (Fig. 2j). Notably, T cells harboring clonotypes shared and expanded across sites localized predominantly to paths 2 and 3 in FA, whereas a subset of CA cells mapped to path 1 (Fig. 2k). Together, FA was associated with a stronger trafficking-dependent systemic T-cell clonal response and preferential representation of CD8^+^ T-cell states with stronger TCR/effector/cytotoxic programs, whereas CA retained greater representation of migratory and IFN-responsive states. Additionally, the myeloid compartment was also profoundly remodeled in FA tumors, with decreased immunosuppressive secreted phosphoprotein 1 (SPP1)-positive macrophages^28^ and concurrently expanded pro-inflammatory CXCL2^+^ subsets (Extended Data Fig. 5), which recruit neutrophils that promote immune responses and T-cell activation.

**Fig. 2.**
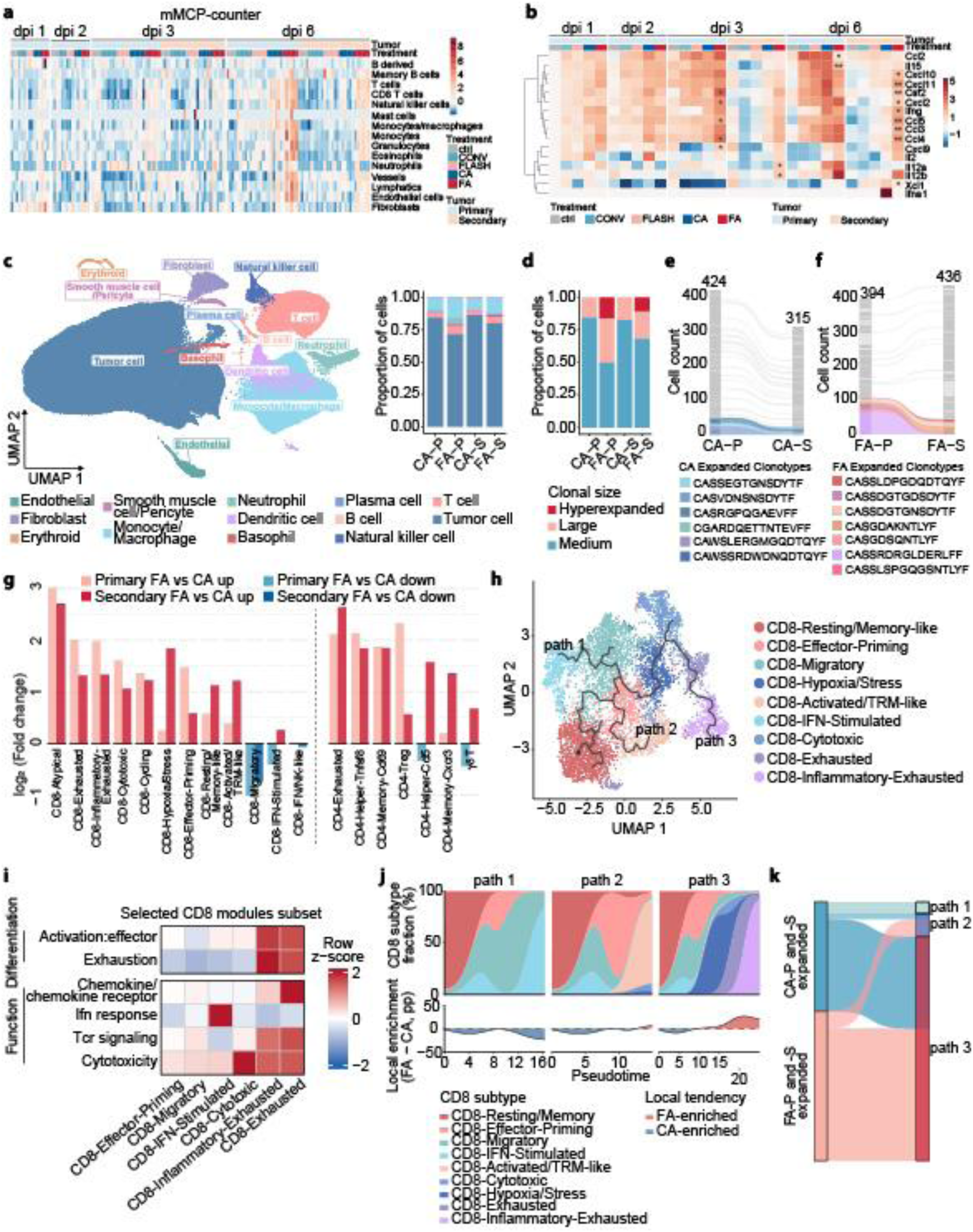
Differential T-cell clonal expansion and trajectory selection between FLASH and CONV X-rays. **a**, Heat map showing immune and stromal cell abundance inferred by mMCP-counter from bulk RNA-seq data. Values are row Z-scores; n = 3–6 biologically independent tumors per group. **b**, Heat map showing relative expression of selected cytokine and chemokine genes, normalized to control tumors at day 1 and displayed as row Z-scores. Asterisks indicate significant differences between CONV X-rays plus αPD-1 (CA) and FLASH X-rays plus αPD-1 (FA) groups (\**P* < 0.05, \*\**P* < 0.01). **c**, Uniform Manifold Approximation and Projection (UMAP) embedding (colored by 13 major cell types) of 228,350 high-quality single cells from primary (P) and secondary (S) tumors collected on days 3 and 6 post-irradiation. The corresponding proportions of these cell types across the indicated groups are shown. n = 2–3 biologically independent tumors per treatment group. **d**, Proportion of hyper, large, and medium-expended TCR clonotypes in T cells. **e**,**f**, Alluvial plots showing TCR clone sharing between irradiated and distant tumors in the CA (**e**) and FA (**f**) groups. Colors denote shared clonotypes expanded at either tumor site. **g**, Fold change in the abundance of 21 T-cell subsets in irradiated and distant tumors in the FA groups versus the CA groups. **h**, Pseudotime trajectory of CD8^+^ T cells showing three major trajectories. **i**, Functional module scores of the major CD8^+^ T-cell subsets retained in the path 1 and path 3 branches. **j**, CD8⁺ T-cell subset composition and treatment-associated distribution along inferred paths. Upper panels show the smoothed proportions of CD8⁺ T-cell subsets along three inferred paths. Lower panels show the relative enrichment of FA versus CA cells from primary tumors along the corresponding paths after balancing the total cell contributions of the two treatment groups. Positive and negative values indicate relative FA and CA enrichment, respectively; the dashed line denotes equal representation of FA and CA cells. Cells located along the shared early segment are included in each corresponding path. pp, percentage points. **k**, Alluvial plot showing the distribution of shared and expanded TCR clonotypes between irradiated and distant tumors across the three CD8^+^ T-cell pseudotime paths in the CA and the FA groups. For **b**, significance was determined by Wilcoxon test. dpi, days post irradiation.

### 3. FLASH X-rays trigger multicellular rewiring toward an antitumor phenotype

GSEA revealed that FA tumor cells consistently exhibited higher enrichment of immune response programs, including IFN signaling and antigen processing and presentation, whereas RNA processing, chromatin remodeling, and cell-cycle programs were generally lower (Fig. 3a and Supplementary Table 1). Consistent with this immune shift, inferred major histocompatibility complex (MHC)-I signaling from malignant cells to CD8^+^ T cells was stronger in FA tumors across all three comparisons, with concordant changes in *H2-D1* and *H2-K1* expression in tumor cells and *Ifng* expression in CD8^+^ T cells (Fig. 3b and Extended Data Fig. 6a–c). Subclustering of malignant cells identified five states, provisionally designated T0–T4. Trajectory analysis aligned T0 (Proliferative), T3 (Immune-reactive), T1 (Intermediate, occupying a position between T3 and T2), and T2 (Injury-like) along a major continuum, whereas the rare T4 (Neural-like) state followed a divergent path (Fig. 3c,d and Extended Data Fig. 6d–h). Proliferative cells declined from day 3 to day 6 post-IR and were consistently lower in FA than CA tumors. Injury-like cells showed the opposite trend, becoming more prominent in FA tumors at day 6 in both irradiated and distant tumors, whereas Intermediate cells remained enriched in CA tumors at the same time (Fig. 3e,f). Thus, the broader immune response program in FA was accompanied by a shift from proliferative to later stress/injury-associated states while retaining an inflammatory context. To characterize these multicellular configurations, we performed CoVarNet^29^ and identified 11 cellular modules (CMs) with distinct treatment- and time-dependent activities (Fig. 3g and Extended Data Fig. 7a,b). At day 3 post-IR, two of three CA primary tumors showed high CM01 activity (dominated by IFN-Stimulated CD8^+^ T cells and γδ T cells), whereas two of three FA tumors showed high CM10 activity (dominated by cytotoxic CD8⁺ T cells). Remaining samples converged to CM08, with CA samples showing intermediate CM01/CM10 activity. By day 6 post-IR, modules diverged further, with FA samples concentrated within a restricted set of dominant modules, whereas CA samples distributed across a more heterogeneous set of CMs. Projection of dominant malignant and CD8^+^ T-cell states of treatment-associated CMs onto their respective trajectories revealed that CA-associated modules were preferentially linked to path 1 CD8^+^ states and centered on the Intermediate malignant state and mid-trajectory CD8^+^ T-cell configurations. In contrast, FA-associated modules more frequently involved path 3 CD8^+^ T-cell states and extended toward later Injury-like malignant states along with later effector/cytotoxic CD8^+^ T-cell states (Fig. 3h and Extended Data Fig. 7c). Multiplex immunofluorescence confirmed enrichment of CD8^+^ISG15^+^ T cells in CA and CD8^+^Granzyme B^+^ T cells in FA, consistent with the IFN-Stimulated and Cytotoxic CD8^+^ components of their respective modules (Fig. 3i). Nearest-neighbor distance analysis further indicated closer proximity of CD4^+^IFITM1^+^ cells to CD8⁺ISG15⁺ T cells in CA but to CD8^+^Granzyme B^+^ T cells in FA (Fig. 3j,k), providing a spatial correlate of the distinct CM configurations. Notably, infiltrating CD8^+^ T cells in CA primary tumors presented a margin-distributed pattern at day 3 post-IR, whereas CD8^+^ T cells in FA tumors penetrated more broadly into the tumor interior (Fig. 3i and Extended Data Fig. 7d), suggesting that a physical barrier restricting T-cell infiltration may contribute to the weaker systemic response in CA.

**Fig. 3.**
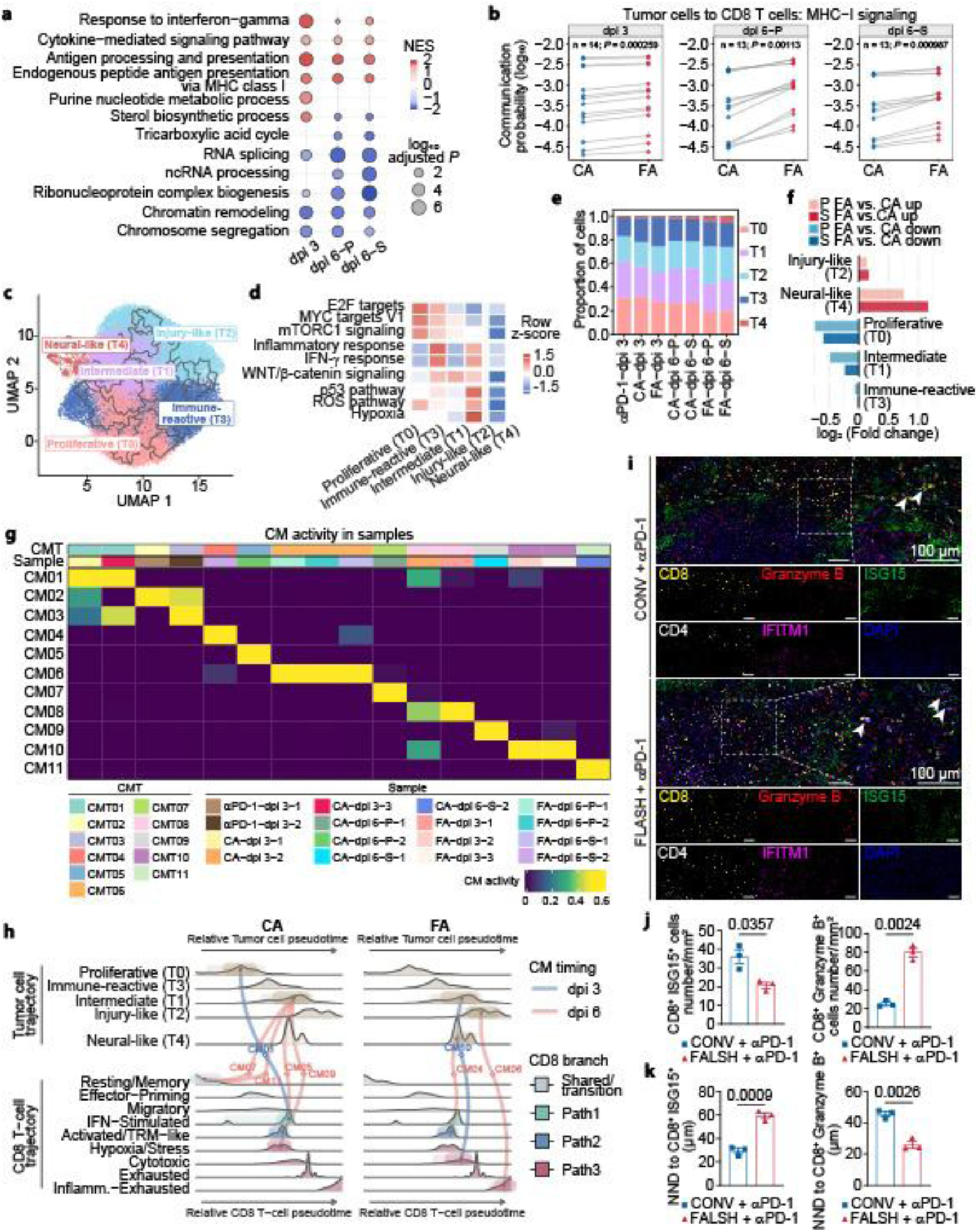
Distinct malignant-cell state dynamics and module activities in FLASH versus CONV tumors. **a**, GSEA bubble plot comparing malignant cells between FA and CA across post-irradiation time points and tumor sites. Positive and negative normalized enrichment score (NES) values indicate enrichment in FA and CA, respectively; color denotes NES and point size denotes –log_10_ (adjusted *P* value). **b**, Inferred MHC-I pathway interactions from tumor cells to CD8^+^ T cells, compared between CA and FA groups; for each group, cells from 2–3 tumors were pooled for analysis, and each point represents one ligand–receptor pair. **c**, UMAP of tumor cells showing five states (T0–T4) with the Monocle3 principal graph. **d**, Heat map of mean UCell scores for selected Hallmark gene sets across the five malignant states (T0–T4). Values are row Z-scores. **e**, Proportions of the five tumor cell states across indicated groups from tumors collected on dpi 3 and dpi 6. **f**, Fold change in the proportions of five tumor cell states in primary (P) and secondary (S) tumors in the FA groups versus the CA groups. **g**, Cellular modules (CMs) activities in scRNA-seq samples from irradiated and distant tumors at dpi 3 and dpi 6 across the αPD-1 alone, and CA and FA groups. Each sample was assigned a cellular module type (CMT) based on its dominant CM and ordered by CMT identity and treatment. **h**, Coupling of tumor states and CD8^+^ T-cell states within treatment-associated CMs. For each CM, the highest-weight tumor states and CD8^+^ T-cell states are positioned according to their relative pseudotime along the corresponding tumor and CD8^+^ T-cell trajectories and connected by a line. Line color denotes CM timing (dpi 3 or dpi 6). CD8^+^ T-cell subset distributions along pseudotime are colored according to trajectory branches. **i**–**k**, Multiplex immunofluorescence assay of irradiated tumors. Representative image (**i**). Scale bar, 100 μm. Arrows indicate adjacent cell pairs. The numbers of CD8⁺Granzyme B⁺ T cells, CD8⁺ISG15⁺ T cells (**j**), and the nearest-neighbor distance (NND) from CD4⁺IFITM1⁺ cells to these subsets (**k**) were quantified and compared between the CA and FA groups. Data are presented as mean ± SEM (n = 3 mice per group). For **d**, significance was determined by the Wilcoxon test. For **j**, statistical significance was determined using two-tailed Student’s *t*-test, and for **k**, using unpaired two-tailed Student’s *t*-test with Welch’s correction. dpi, days post irradiation.

### 4. FLASH X-rays reverse NET-mediated CD8⁺ T-cell exclusion and concomitant immunosuppression

Radiotherapy can induce intratumoral barriers, particularly CAF-mediated ones, that restrict CD8⁺ T-cell infiltration and impair abscopal responses^8,30^. However, alpha-smooth muscle actin (α-SMA), a critical marker of activated CAFs, did not differ between the FA and CA groups (Extended Data Fig. 8a–c). We therefore asked whether neutrophil extracellular traps (NETs), physical barriers formed during NETosis^11,31^, blocked CD8^+^ T-cell entry into the tumor center. In contrast to FA tumors, primary CA tumors displayed a strong fluorescent strip-like pattern positive for both citrullinated histone H3 (Cit H3) and myeloperoxidase (MPO) at day 3 post-IR, indicative of NET formation (Fig. 4a,b). Consistently, CD8⁺ T cells in the CA group were largely confined to the tumor margin, aligning with the NET-rich strips, whereas they were evenly distributed throughout the tumors in the FA group, suggesting that NETs impeded CD8^+^ T-cell infiltration into the irradiated tumor center (Fig. 4a–c). *In vivo* CD8^+^ T-cell depletion abrogated the clearance of both primary and secondary tumors and eliminated the survival benefit (Extended Data Fig. 8d–k), underscoring their essential role in the abscopal response. To test whether disrupting NETs could restore T-cell infiltration and abscopal efficacy, we used GSK484, a NET-specific inhibitor (Fig. 4d). GSK484 largely dissolved the characteristic cord-like NETs in CA-treated primary tumors (Fig. 4e,f) and increased tumor-center CD8^+^ T cells from < 15% to ∼ 40% (*P* = 0.0002) (Fig. 4g). This enhanced T-cell infiltration translated into a marked survival benefit, from 16.6% to 64.3% (*P* = 0.0072) (Fig. 4h), suggesting systemic immune reactivation. Although GSK484 was ineffective as a monotherapy or with CONV IR or αPD-1, it synergized with CA treatment to rescue metastatic clearance from 2/12 to 9/14 (Fig. 4i,j) while sustaining primary tumor response. Tumor rechallenge experiments further confirmed the durability and specificity of the restored immunity (Fig. 4k). Mice cured by CA + GSK484 rejected MC38 rechallenge 90 days later, but not the unrelated B16-F10 tumors (Fig. 4l,m), indicating antigen-specific immunological memory. This pattern mirrored the durable antitumor immunity elicited by FA, reinforcing that NET inhibition is a key mechanism underlying the superior systemic response to FLASH-based radioimmunotherapy.

**Fig. 4.**
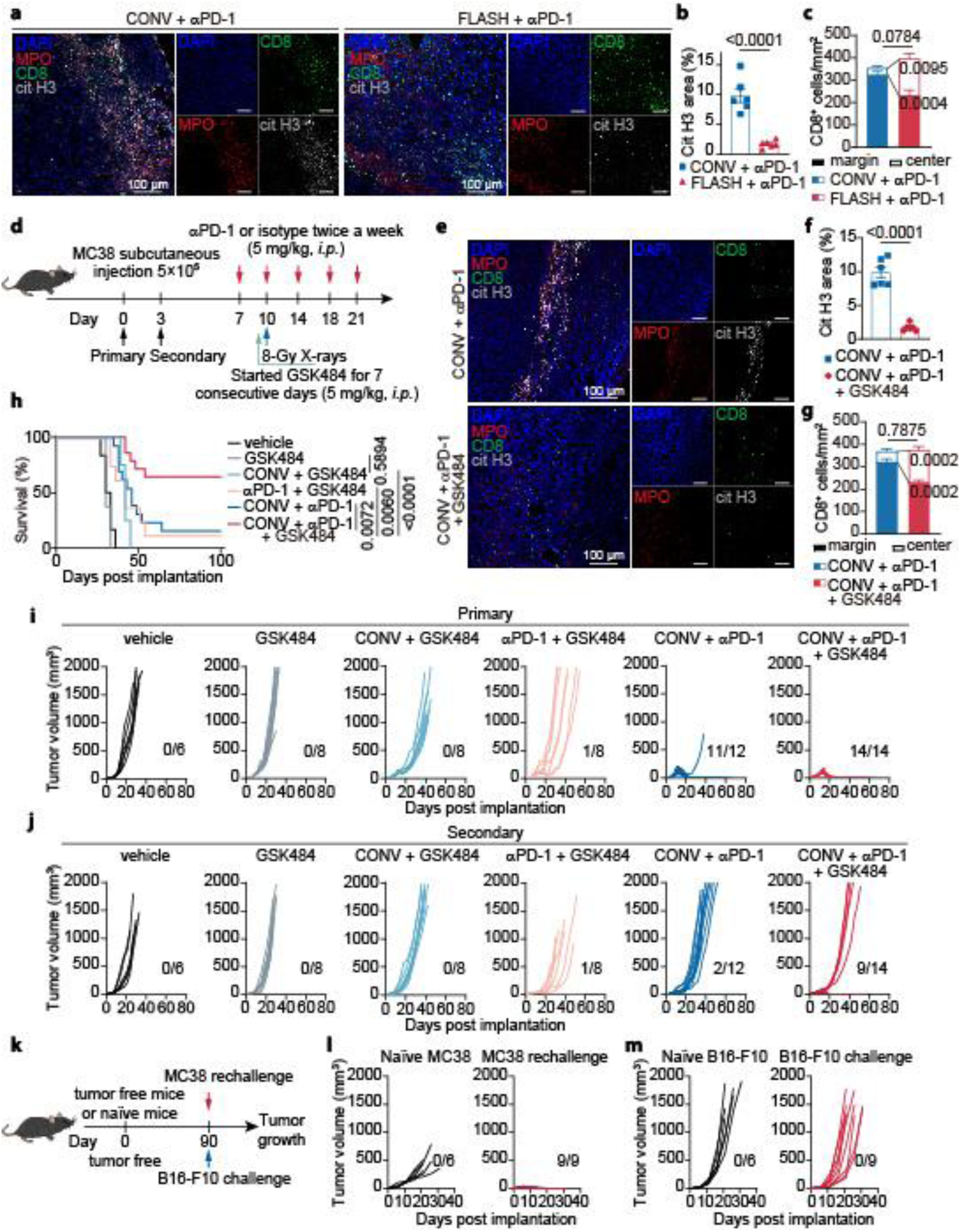
Reversed NET-mediated CD8+ T-cell exclusion by FLASH versus CONV X-rays. **a**–**c,** The presence of NETs (Cit H3, citrullinated histone H3, gray; MPO, red) and the spatial distribution of CD8⁺ T cells (green) within irradiated tumors were examined at 3 days post-irradiation. Representative confocal images are shown in (**a**) (20× objective; scale bar, 100 μm). Quantifications of NETs (percentage of Cit H3-positive area) and CD8⁺ T-cell infiltration (CD8⁺ cells/mm²) are presented in (**b**) and (**c**), respectively (n = 6 mice per group). **d**, Schema for evaluating the role of GSK484 in inhibiting NET formation and modulating the abscopal effect induced by CONV X-rays. **e**–**g**, The presence of NETs and the distribution of CD8⁺ T cells within irradiated tumors from the CA + vehicle group or the CA + GSK484 group were examined at 3 days post-irradiation. Representative confocal images are shown in (**e**) (20× objective; scale bar, 100 μm). Quantifications of NETs and CD8⁺ T-cell infiltration are presented in (**f**) and (**g**), respectively (n = 6 mice per group). **h**–**j**, Mouse survival and tumor growth are evaluated in the indicated groups. Kaplan–Meier survival curves are shown in (**h**), primary (**i**) and secondary (**j**) tumor volumes of individual mice are plotted, and the ratios (tumor-free mice/total mice) are indicated for each treatment group. **k**, Schema for evaluating the antitumor immune memory of tumor-free mice from the CA + GSK484 group. **l**,**m**, Tumor growth curves of naïve (black) mice and tumor-free (red) mice (previously treated with CA + GSK484) after rechallenge with MC38 (**l**) or with the unrelated B16-F10 melanoma (**m**), and the ratios (tumor-free mice/total mice) are indicated for each treatment group. Data are presented as mean ± SEM. Statistical significance was assessed using unpaired two-tailed Student’s *t*-test (**b**,**c**,**f**,**g**) and log-rank test (**h**).

### 5. FLASH X-rays inhibit NET formation by sustaining chromatin compaction

FLASH IR may instantaneously deplete oxygen and limit reactive oxygen species (ROS) generation^16,32^, thereby partially suppressing NETosis^33^. However, ROS levels in *ex vivo*-cultured mouse bone marrow neutrophils were comparable 30 min post-IR at different dose rates (Extended Data Fig. 9a,b). Moreover, conditioned medium from MC38 cells irradiated with CONV or FLASH X-rays induced comparably minimal NET formation in neutrophils (Extended Data Fig. 9c,d). In contrast, direct 8-Gy CONV IR induced markedly more NETs than direct FLASH X-rays (Fig. 5a,b), indicating that FLASH X-rays acted directly on neutrophils rather than indirectly via the tumor microenvironment. Although macrophages can also form similar extracellular DNA scaffolds, termed macrophage extracellular traps (METs)^34^, Cit H3 and PAD4 levels in mouse bone marrow-derived macrophages (BMDMs) remained unchanged after CONV IR (Extended Data Fig. 9e–j), thereby ruling out a macrophage origin of the extracellular DNA scaffolds. We previously proposed that the FLASH sparing effect arises from chromatin compaction during dose delivery (“DNA integrity” hypothesis)^25^. In line with this, the extremely short FLASH delivery time prevented chromatin decondensation and abrogated the global histone H3 acetylation (H3ac) and the K9/K27 acetylation that were rapidly upregulated after CONV IR (Fig. 5c,d). Key enzymes such as PARP1 anchor to DNA break sites within seconds after radiation-induced DNA damage, catalyzing both autoPARylation and PARylation of histones^35,36^, driving early chromatin relaxation and subsequent protein acetylation^37^. Consistent with this mechanism, FLASH X-rays dramatically reduced CONV-induced protein PARylation in neutrophils (Fig. 5e). PJ34, a reversible, highly selective inhibitor of PARP1/2 enzymatic activity^38^, abrogated CONV-induced PARylation, reversed H3 acetylation, and blocked NET formation *in vitro* (Fig. 5e–h), indicating that chromatin PARylation drives the excessive chromatin decondensation during CONV dose deposition. To dissect whether the differential effects arose from the instantaneous dose delivery or the ultradense pattern, we split a single 8-Gy FLASH giant pulse into two 4-Gy pulses separated by a 5-min interval (Extended Data Fig. 9k). This approach deliberately extended the total delivery time while preserving the ultradense mode and macro-pulse dose rate. This split-dose regimen recapitulated CONV-triggered PARylation (Fig. 5i), H3 acetylation (Fig. 5j), and NET formation (Fig. 5k), all of which were reversed by PJ34 (Fig. 5i,j and Extended Data Fig. 9l). Thus, the ultrashort dose-delivery duration, rather than the ultradense pattern, determines the chromatin modification cascade that drives NETosis.

**Fig. 5.**
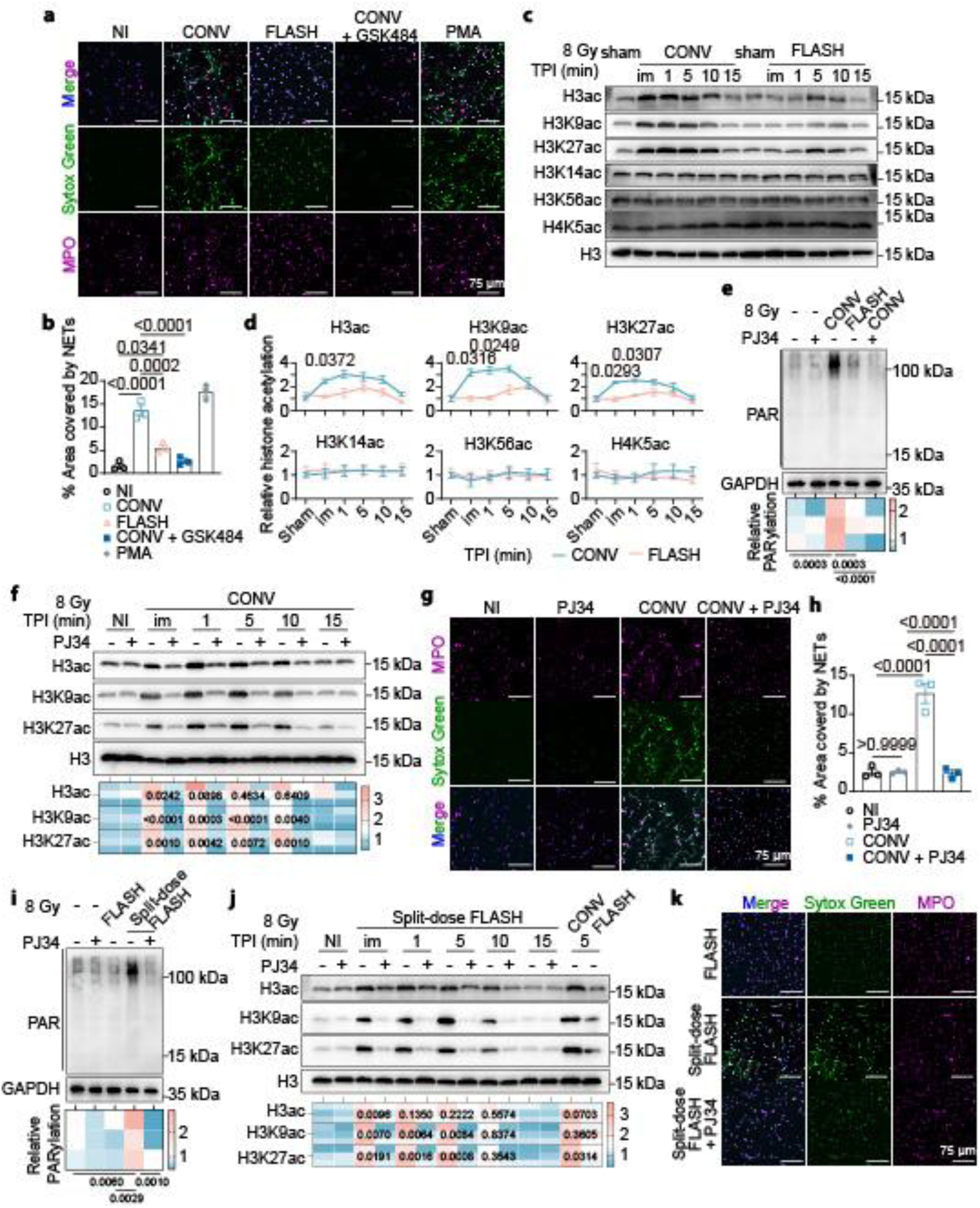
Chromatin compaction in neutrophils correlates with reduced NETs under FLASH. **a**,**b**, NETs were visualized by staining for MPO (magenta) and extruded DNA (SYTOX Green) at 4 h after the indicated treatments. Representative confocal images are shown in (**a**), and quantification of NETs is shown in (**b**). **c**,**d**, Western blot analysis of histone acetylation levels in mouse neutrophils following 8-Gy CONV or FLASH irradiation at the indicated times post-irradiation (**c**) and the corresponding quantification (**d**). **e**–**h**, Radiation-induced PARylation was inhibited by PJ34 pretreatment. Protein PARylation levels (**e**) and histone acetylation levels (**f**) were examined and quantified by Western blot immediately after 8-Gy irradiation. Representative confocal images of NETs at 4 h after 8-Gy CONV irradiation are shown in (**g**), and quantification of NETs is shown in (**h**). **i**–**k**, FLASH X-ray exposure was fractionated equally by introducing a 5-min interval (termed split-dose FLASH). Protein PARylation levels (**i**) and histone acetylation levels (**j**) were examined and quantified by Western blot immediately after FLASH irradiation (either continuous or fractionated) with or without PJ34 pretreatment, and NET formation was assessed by confocal microscopy at 4 h after 8-Gy FLASH irradiation (**k**). Data are presented as mean ± SEM. Statistical significance was assessed using one-way ANOVA (**b**,**e**,**f**,**h**,**i**,**j**) and two-way ANOVA followed by Šídák’s multiple comparisons test (for each time point between groups) (**d**). NI, Non-irradiated; TPI, time post-irradiation; im, immediately.

### 6. PARP inhibition eliminates NETs and enhances the abscopal effect under radioimmunotherapy

To validate whether split-dose FLASH X-rays trigger NETosis *in vivo* (Fig. 6a), we sampled irradiated tumors at day 3 post-IR for MPO and Cit H3 colocalization. As shown in Fig. 6b, split-dose FLASH + αPD-1 recapitulated the NET-rich barrier seen in CA-treated primary tumors, excluding CD8⁺ T cells from the tumor center. Indeed, central CD8⁺ T-cell density dropped from 162 ± 23 cells/mm² in the FA group to 51 ± 7 cells/mm² (*P* = 0.0035) (Fig. 6c,d). Consistently, split-dose FLASH reduced the abscopal effect from 60% to 30% without compromising primary tumor control (Fig. 6e), and this impaired systemic immunity drove survival down to levels comparable to the CA group (Fig. 6f). Given that chromatin modifications before dose completion drove NETosis and abrogated the abscopal response, we asked whether CA-treated mice could be rescued by pharmacologically suppressing chromatin modifications during IR. To this end, we administered PJ34 or the investigational PARP1/2 inhibitor veliparib intraperitoneally one day before IR and daily thereafter for 6 days (Fig. 6g). Notably, PARP inhibition nearly abolished CONV-induced NETs, as evidenced by loss of Cit H3 and MPO co-localization (Fig. 6h,i), and restored CD8⁺ T-cell central infiltration (Fig. 6j). Importantly, while PJ34 did not improve overall survival or tumor control rates in the CONV-alone group, it significantly prolonged survival in the CA group (from 15.4% to 54.5%) and enhanced metastatic lesion clearance from 2/13 to 6/11 (Fig. 6k,l), approaching those of the FA group. Together, these findings establish that neutrophil chromatin preservation, achieved either through ultrashort dose delivery or through pharmacological PARP inhibition, suppresses NETosis and unleashes systemic antitumor immunity (Extended Data Fig. 10).

**Fig. 6.**
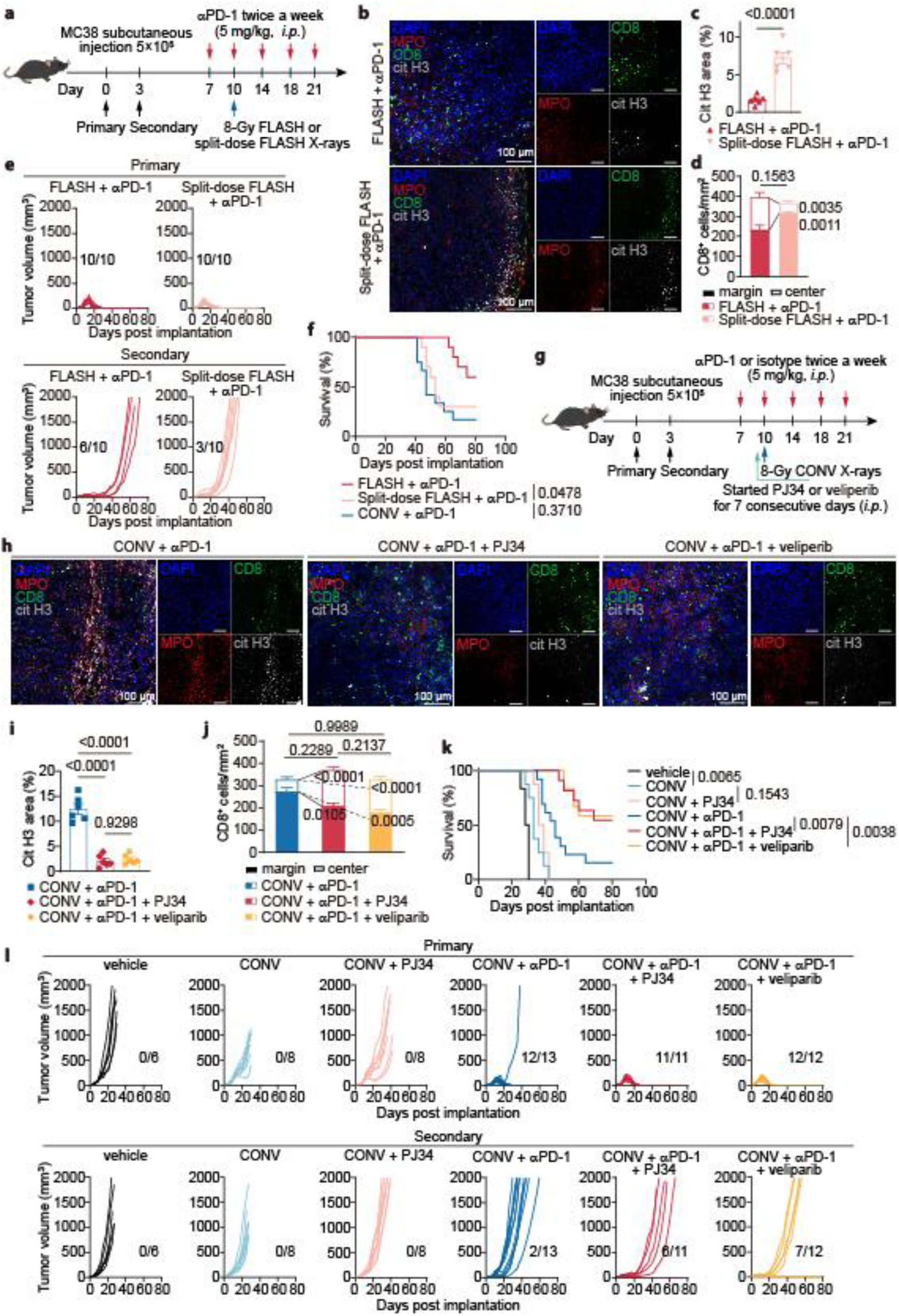
PARP inhibition suppresses NETs and restores abscopal responses under radioimmunotherapy. **a**, Schema for evaluating the effect of introducing a 5-min interval during FLASH irradiation (split-dose FLASH) on NET formation within irradiated tumors and the abscopal effect induced by FLASH combined with αPD-1. **b**–**d**, The presence of NETs and the distribution of CD8⁺ T cells within irradiated tumors from the FLASH + αPD-1 group or the split-dose FLASH + αPD-1 group were examined at day 3 post-irradiation. Representative confocal images are shown in (**b**) (20× objective; scale bar, 100 μm). Quantifications of NETs and CD8⁺ T-cell infiltration are presented in (**c**) and (**d**), respectively (n = 6 mice per group). **e**,**f**, Survival and tumor growth of mice from the continuous or spilt-does FLASH plus αPD-1 group were evaluated. Kaplan–Meier survival curves are shown in (**e**), primary and secondary tumor volumes of individual mice are plotted (**f**), and the ratios (tumor-free mice/total mice) are indicated for each treatment group. **g**, Experimental design for evaluating the effect of PJ34- or veliparib-mediated PARP inhibition on NET formation within irradiated tumors and the abscopal effect induced by CONV combined with αPD-1. **h**–**j**, The distribution of NETs and CD8⁺ T cells in irradiated tumors from the CA + vehicle, CA + PJ34, and CA + veliparib groups was evaluated on day 3 post-irradiation. Representative confocal images (20× objective; scale bar, 100 μm) are shown in (**h**), with corresponding quantifications of NETs (**i**) and CD8⁺ T-cell infiltration (**j**) (n = 6 mice per group). **k**,**l**, Survival and tumor growth of mice treated with CONV plus αPD-1 with or without a PARP inhibitor were evaluated. Kaplan–Meier survival curves are shown in (**k**), individual primary and secondary tumor volumes are plotted in (**l**), and the tumor-free/total mice ratios are indicated for each group. Data are shown as mean ± SEM. Statistical significance was assessed using unpaired two-tailed Student’s *t*-test (**c**,**d**), one-way ANOVA (**i**,**j**) and log-rank test (**f**,**k**).

## Discussion

Decades of technical refinement have improved radiotherapy’s spatial precision, yet single-cell resolution remains out of reach^39^, leaving intratumoral immune cells inevitably exposed to radiation damage^7^. FLASH radiotherapy, by radically compressing dose delivery into milliseconds^12,15^, offers a temporal solution to this spatial limitation^24^. Our study establishes that the therapeutic advantage of FLASH over CONV IR in combination with αPD-1 lies not simply in its ultrahigh pulse dose rate, but in the ultrashort temporal continuity of dose delivery, which preserves chromatin compaction in intratumoral neutrophils and prevents a cascade of pro-NETotic chromatin modifications that would otherwise abrogate systemic antitumor immunity.

Mechanistically, we show that CONV IR, owing to its prolonged delivery time, allows PARP1 to anchor to DNA breaks and catalyze histone PARylation within seconds, triggering rapid chromatin decondensation and histone H3 acetylation. The aforementioned PARylation occurring before dose completion predisposes neutrophils to NETosis, a process we confirmed by split-dose experiments in which introducing a 5-min pause into FLASH delivery to mimic the CONV dose-delivery time recapitulated the CONV phenotype. Thus, the ultrashort dose-delivery duration of FLASH effectively “outruns” the PARylation-chromatin relaxation axis, maintaining chromatin in a compact state and blocking NET formation. This mechanistic insight extends our previously proposed the “DNA integrity” hypothesis, which posited that FLASH mitigates toxicity by reducing cytosolic DNA fragments derived from chromatin relaxation^25^, to one that specifically protects neutrophil chromatin from IR-induced aberrant modification.

Functionally, NETs generated under CONV IR form physical barriers that exclude CD8^+^ T cells from the tumor core, a phenomenon we visualized as NET-rich strips at the tumor margin. Targeted disruption of NETs with GSK484 restored T-cell infiltration and rescued the abscopal response, mirroring the effect of FLASH. Importantly, FLASH not only increased the quantity of tumor-infiltrating CD8^+^ T cells but also improved their quality: TCR sequencing revealed that FLASH promoted cross-tumor clonal sharing and expansion, and pseudotime trajectory analysis showed that shared clones preferentially differentiated into cytotoxic/effector-memory paths (path 2 and 3), whereas CONV skewed them toward an IFN-stimulated, low-cytotoxicity state (path 1). Thus, by preserving neutrophil homeostasis, FLASH licenses a more robust and durable T-cell response, with superior systemic immune surveillance against metastases.

The translational implications of our findings are twofold. First, our split-dose results pose a formidable challenge to clinical implementation of FLASH, particularly in multi-beam delivery using a rotatable gantry, where unavoidable inter-field pauses could compromise the FLASH effect^40,41^. This calls for technological solutions that maintain beam continuity (*e.g.*, using a pluridirectional multi-source linac array^42^) or redefine treatment planning to minimize interruptions. Second, we demonstrate that pharmacological inhibition of PARylation, using PJ34 or veliparib, combined with CA treatment mirrors the immune-sparing benefits of the FA regimen, improving survival and clearance of metastatic lesions. Notably, this effect was independent of direct DNA repair inhibition in tumor cells, revealing a non-canonical, immune-centric mechanism that positions PARP inhibitors as a near-term translatable strategy to enhance radioimmunotherapy even where FLASH equipment is unavailable. Given the ongoing clinical trials combining PARP inhibitors with radiotherapy and immunotherapy (e.g., NCT04837209, NCT04711824, NCT04380636, NCT05411094), our data provide a mechanistic rationale for patient selection and treatment sequencing.

We acknowledge several limitations. The subcutaneous MC38 model, while adequate for dissecting immune mechanisms, does not fully recapitulate the complex TIME of spontaneous or orthotopic tumors. Optimal FA regimens, such as FLASH parameters, may vary with tumor type or immune context. Additionally, the long-term safety of combining PARP inhibitors with high-dose radiotherapy warrants further investigation. Nonetheless, our study establishes a mechanistic connection between dose-delivery kinetics, neutrophil chromatin biology, and systemic antitumor immunity, and offers a dual strategy—either via FLASH technology or pharmacological PARP inhibition — to broaden the therapeutic window of radioimmunotherapy in both directions.

In summary, by extending the “DNA integrity” hypothesis to encompass neutrophil-specific chromatin protection, we redefine the FLASH effect as not merely normal-tissue sparing, but also immune-sparing and immunopotentiating. Our findings pave the way for clinical strategies that harness either physical (FLASH) or pharmacological (PARP inhibitors) means to preserve chromatin stability in neutrophils, thereby unleashing the full potential of radioimmunotherapy against metastatic disease.

## Methods

### Cell lines, mice and ethics statement

Mouse colon carcinoma MC38 cells and B16-F10 cells, both purchased from Shanghai FuHeng Biotechnology Co., Ltd., were cultured in RPMI-1640 medium (for MC38) or high-glucose DMEM (for B16-F10), supplemented with 10% fetal bovine serum, penicillin (100 U/mL), and streptomycin (100 μg/mL) at 37 °C in a 5% CO₂ atmosphere. No additional authentication was performed beyond the identity provided by the supplier. Mycoplasma testing was negative. All experiments were performed with cells within passages 3–10 after thawing. Specific pathogen-free (SPF) male C57BL/ 6J mice (6–8 weeks old) were obtained from GemPharmatech Co., Ltd. (Chengdu, China). Mice were housed under a 12 h/12 h light–dark cycle at 24 °C with ≤ 5 mice per cage. The study was conducted in compliance with local animal welfare laws and policies. All animal experimental procedures were approved by the Animal Welfare and Ethics Committees of Fudan University (Approval No. 2021JS-FYS-001) and Soochow University (Approval No. SUDA20240911A33) and complied with the ARRIVE 2.0 guidelines. Tribromoethanol anesthesia was used for irradiation procedures.

### Irradiator and dosimetry

X-ray FLASH irradiation experiments were performed using a compact single high-energy X-ray source (CHEXS) device (10 MV)^43^ at a mean dose rate of approximately 100 Gy/s, whereas conventional dose-rate (CONV) irradiation at 0.03 Gy/s was delivered using a clinically applied 6 MV Elekta Precise linac (Elekta AB). The percent depth dose (PDD) curve was consistent with the previous description^43^. For *in vivo* studies, mice were anesthetized and positioned prone on a 20 × 20 cm PMMA fixation plate with a thickness of 2 mm for vertical irradiation. For *in vitro* studies, cells in T12.5 culture flasks filled with pre-warmed medium were placed horizontally for irradiation.

A fast current transformer (FCT; Bergoz ICT-082-012-05) was installed at the entrance of the achromatic beam transport section to monitor electron beam intensity. Real-time monitoring of X-ray dose and spatial distribution during irradiation was performed using a custom-built low-pressure ionization chamber. An 8-cm-thick lead secondary collimator, with aperture sizes of 2 × 2 cm (for tumor irradiation) and 8 × 5 cm (for cell irradiation), was used to define the FLASH irradiation field. The secondary collimator was equipped with a flattening filter to improve dose uniformity, and a PMMA tissue compensator served as a build-up material.

Dose verification was performed using EBT4 films (Ashland Inc., Covington, KY, USA) and a PTW diamond detector (model 60025). EBT4 films were positioned in a solid water phantom at a depth of 0.5 cm, corresponding to the tumor depth, to measure the absolute dose at the beam center, the mean dose within 80% of the field size, and the dose distribution profiles. The PTW diamond detector was placed at the same depth to measure the absolute dose at the beam center, which was cross-referenced with the central dose obtained from EBT4 films. The flatness of both the FLASH and CONV irradiation fields—defined as the ratio of the maximum dose to the mean dose within 80% of the field size—was approximately 1.05 in both the horizontal and vertical directions within the 2 × 2 cm field. The total irradiation dose was controlled by adjusting the exposure time, and the dose error for each mouse was within ± 1%. These flatness data were consistent with previously reported values^44^.

### Animal model and treatment

Mice (6–8 weeks old) were subcutaneously injected in the left flank with 100 μL of cell suspension containing 5 × 10⁵ MC38 cells on day 0 to establish the primary tumor. Three days later, the same number of cells was injected into the contralateral flank to generate the secondary tumor. When the primary tumor volume reached approximately 100 mm³ (on day 10), it was locally irradiated with a single dose of 8-Gy delivered at either a conventional dose rate (CONV) or an ultrahigh dose rate (FLASH). Before irradiation, mice were anesthetized with 250 mg/kg tribromoethanol for positioning. On days 7, 10, 14, 18, and 21 after primary tumor inoculation, mice received intraperitoneal injections of anti-PD-1 antibody (Bio X Cell, BE0146) or isotype control antibody (Bio X Cell, BE0089) at 5 mg/kg. Tumor volumes were evaluated every two to three days with a digital caliper and were calculated using the following formula (V = longest dimension × perpendicular dimension × perpendicular dimension × 0.5) as previously described^6^. Mice were euthanized by CO₂ asphyxiation when the tumor diameter exceeded 20 mm or when tumors became ulcerated. For FTY720 treatment, mice received a loading dose of 25 μg of FTY720 (Selleck, S5002) per mouse via intraperitoneal injection one day before irradiation, followed by daily maintenance doses of 5 μg/mouse/day until the predefined endpoints^45^. For CD8⁺ T-cell depletion, mice received intraperitoneal injections of αCD8 (Selleck, A2102) at 200 μg per injection, starting one day before irradiation, twice weekly, for a total of five injections. To inhibit NET formation, GSK484 hydrochloride (TargetMol, TQ0067) dissolved in ddH_2_O was intraperitoneally injected into tumor-bearing mice at 5 mg/kg once daily for 7 days, starting one day before irradiation. To inhibit PARP-mediated PARylation, PJ34 HCl (Selleck, S7300, 5 mg/kg) or veliparib dihydrochloride (Selleck, E7366, 10 mg/kg) dissolved in ddH_2_O was intraperitoneally injected into tumor-bearing mice once daily for 7 days, starting one day before irradiation.

### Clonogenic Survival Assay

Cultured MC38 cells at 80–90% confluence were irradiated with CONV or FLASH X-rays at doses of 2, 4, 6, or 8 Gy. Immediately after irradiation, cells were trypsinized, resuspended in fresh complete medium, and counted. Appropriate numbers of cells were seeded into 6-well plates in triplicate per dose: 300 cells (0 Gy), 500 (2 Gy), 3,000 (4 Gy), 10,000 (6 Gy), and 80,000 (8 Gy), and then cultured for colony formation. Cells were cultured for 12 days, fixed with 4% paraformaldehyde for 15 min, and stained with 0.5% crystal violet for 30 min. Colonies (≥ 50 cells) were counted. Survival curves were fitted by nonlinear regression using the multi-target single-hit model.

### Quantitative Real-Time Polymerase Chain Reaction

RNA was extracted from MC38 cells using TRIzol (Invitrogen, 15596018CN) and reverse-transcribed to cDNA with FastKing gDNA Dispelling RT SuperMix (TIANGEN, KR118) according to the manufacturer’s instructions. cDNA samples were further amplified using a fast real-time PCR system (Bio-Rad Laboratories, CFX96™) with the SYBR Green PCR kit (TIANGEN, FP217) according to the manufacturer’s instructions. Relative mRNA expression was calculated using the 2^–ΔΔCt method normalized to *Actb*. The primers used were as follows: *H2-K1* forward, 5′-CAGGTGGAAAAGGAGGGGAC-3′, *H2-K1* reverse, 5′-CTGAGGGCTCTGGATGTCAC-3′; *Cd274* forward, 5′-CCACATTTCTCCACATCTAGCA-3′, *Cd274* reverse, 5′-TCCATCCTGTTGTTCCTCATTG-3′; *Hmgb1* forward, 5′-TCCAGAGATGGCGCACTTAT-3′, *Hmgb1* reverse, 5′-TGCTTGTCATCTGCTGGCAG-3′; *Actb* forward, 5′-GTGACGTTGACATCCGTAAAGA-3′, *Actb* reverse, 5′-GCCGGACTCATCGTACTCC-3′.

### Tumor rechallenge experiment

For tumor rechallenge experiments, 90 days after having rejected their primary tumor challenges, tumor-free mice were rechallenged by subcutaneous implantation of 5 × 10⁵ MC38 cells into the right flank and 2.5 × 10⁵ B16-F10 cells into the left flank. Age-matched naïve mice received the same tumor implantations as controls.

### Co-culture assay for splenocyte activation

Splenocytes were isolated from tumor-free mice 90 days after complete tumor regression and from age-matched naïve mice, then co-cultured with pre-plated MC38 or B16-F10 tumor cells at a 10:1 effector-to-target ratio for 24 h. Cells were then harvested for flow cytometry to analyze activation markers (CD69 and IFN-γ) in CD4⁺ and CD8⁺ T cells.

### Bulk RNA-seq

Total RNA was extracted from tumor tissues using a DNA/RNA/Protein Kit (Omega, R6734-01) according to the manufacturer’s instructions. RNA quality was assessed by agarose gel electrophoresis, NanoDrop spectrophotometry and Agilent 2100 Bioanalyzer analysis. Qualified RNA samples were used for poly(A)+ mRNA enrichment with oligo(dT) magnetic beads, followed by cDNA synthesis, end repair, A-tailing, adaptor ligation, purification, PCR amplification and circularization. Libraries were quantified using Qubit 3.0, insert sizes were assessed using the Agilent 2100 Bioanalyzer and effective library concentrations were determined by qPCR. DNA nanoballs were generated from qualified libraries and sequenced on an MGI-2000 platform to generate 150 bp paired-end reads.

### Bulk RNA-seq processing and mMCP-counter analyses

Raw sequencing reads were processed with SOAPnuke (v1.5.6) to remove adaptor sequences, low-quality reads and reads containing excessive ambiguous bases. Filtered paired-end reads were aligned using Bowtie2 (v2.2.5)^46^ to an RSEM-compatible transcriptome reference generated from UCSC mm9 RefSeq gene models (mm9_refGene.txt). Gene-level FPKM values were estimated from the resulting BAM files using RSEM (v1.2.31)^47^ in paired-end, non-strand-specific mode (--forward-prob 0.5).

Cell-type abundance scores were calculated using mMCP-counter (v1.1.0)^48^. Entrez gene identifiers in the FPKM matrix were mapped to mouse Ensembl gene IDs and official gene symbols using the bitr function in clusterProfiler (v4.6.2)^49^ with org.Mm.eg.db as the annotation database. Unannotated genes and non-unique gene-symbol mappings were removed. The resulting matrix was transformed as log_2_ (FPKM + 1) and analyzed using the GRCm38 mouse signatures implemented in mMCP-counter. Scores were interpreted as relative abundance estimates and compared across samples within each cell type.

### Single-cell RNA and V(D)J (TCR) library preparation and sequencing

Single-cell libraries were prepared using two workflows according to the sampling time after radiotherapy. Samples collected on day 3 post-irradiation (dpi 3) were processed for single-cell 3′ gene expression (GEX) profiling, whereas samples collected on day 6 post-irradiation (dpi 6) were subjected to integrated single-cell 5′ GEX and TCR V(D)J sequencing.

For 5′ GEX and TCR V(D)J libraries, library preparation was performed using the DNBelab C4 Single-Cell 5′ RNA & V(D)J Library Preparation Kit (MGI, 940-001988-00). Briefly, single-cell suspensions were co-partitioned with barcoded magnetic beads and oil-phase reagents on a DNBelab C-TaiM 4 System chip to generate water-in-oil droplets for single-cell encapsulation and barcoding. After cell lysis, poly(A) RNA was captured by oligo(dT) and template-switching oligonucleotide (TSO) sequences on the bead surface. Full-length cDNA containing cell barcodes and unique molecular identifiers (UMIs) was generated via template-switching reverse transcription, followed by cDNA amplification. Amplified cDNA was then fragmented and split for construction of 5′ GEX libraries and TCR V(D)J libraries. TCR libraries were enriched using two rounds of nested PCR: the first-round PCR employed outer primers targeting TCR constant regions to enrich TCR fragments, and the second-round PCR used inner primers to further amplify products and incorporate sequencing adapters.

For dpi 3 samples, 3′ GEX libraries were prepared using the DNBelab C4 3′ Single-Cell RNA Library Preparation Kit (MGI, 940-001818-00) according to the manufacturer’s instructions, including droplet-based single-cell encapsulation, mRNA capture, reverse transcription, cDNA amplification, and library purification.

All libraries passing quality control were quantified and sequenced on the DNBSEQ-T7 platform in paired-end mode. Raw sequencing data were processed using DNBC4Tools (v2.1.3) for cell calling and gene expression quantification, and TCR clonotypes were additionally assembled and analyzed for the 5′ GEX + TCR V(D)J libraries. All reads were aligned to the mouse reference genome GRCm39.

### scRNA-seq data processing

All single-cell gene expression data were processed and quality-controlled using Scanpy (v1.9.3)^50^. Potential doublets were identified and removed on a per-sample basis using the sc.pp.scrublet function.

Given the different count-depth characteristics of the 3′ and 5′ chemistries, distinct filtering thresholds were applied. For the 5′ datasets, cells with fewer than 100 or more than 6,000 detected genes, or with total UMI counts (total_counts) greater than 30,000, were excluded. For the 3′ datasets, cells with fewer than 100 or more than 8,000 detected genes, or with total_counts greater than 80,000, were removed. In addition, for both 3′ and 5′ datasets, cells with a mitochondrial transcript fraction exceeding 5% were filtered out. After quality control, the retained 3′ and 5′ datasets were merged for subsequent integrated downstream analyses.

### Clustering and determination of the major cell types

After preprocessing, the integrated 3′ and 5′ datasets were normalized and log-transformed (log1p). Highly variable genes (HVGs) were identified separately for the 3′ and 5′ datasets (n = 1,500 each), and the union of the two HVG sets was used as the global feature set. Expression values of the selected features were then scaled.

Dimensionality reduction was performed by principal component analysis (PCA). Batch effects across datasets and individual samples were corrected using Harmony implemented in Scanpy external (sc.external.pp.harmony_integrate). The Harmony-corrected principal components were used to construct the k-nearest-neighbor (kNN) graph. Uniform Manifold Approximation and Projection (UMAP) was applied for visualization, and unsupervised clustering was carried out using the Leiden algorithm.

Cluster marker genes were identified using sc.tl.rank_genes_groups with the Wilcoxon rank-sum test. Major cell types were annotated by comparing cluster marker genes with canonical cell-type markers curated from published literature and public databases^51,52^. For downstream fine-grained analyses, tumor cells were subclustered using the Scanpy workflow, whereas other major cell types were subclustered and further annotated using the Seurat (v5.1.0) pipeline^53^.

For compositional analyses, malignant-cell-state proportions were calculated relative to the total malignant-cell population within each group, whereas proportions of other cell types and states were calculated relative to the total number of cells in each group.

### inferCNV analysis

To support the malignant-cell annotation, large-scale copy number variation (CNV) profiles were inferred using inferCNV (v1.22.0). Genes were ordered according to their genomic positions in the mouse GRCm39 reference genome, and B and T cells were used as the normal reference population. inferCNV was run with cutoff = 0.1 and cluster_by_groups = TRUE, with denoising and hidden Markov model (HMM) prediction disabled.

### Differential expression and functional enrichment analyses

Functional enrichment analyses were performed using the R package clusterProfiler (v4.6.2)^49^ with default parameters unless otherwise specified. For gene set enrichment analysis (GSEA), differential expression between experimental groups was assessed using the FindMarkers function in Seurat with logfc.threshold = 0 and min.pct = 0. Genes were ranked by log_2_ fold change (log2FC), and GSEA was performed using the GSEA function. For Gene Ontology (GO) enrichment analysis of malignant cell states, differentially expressed genes were identified using the sc.tl.rank_genes_groups function in Scanpy. Upregulated genes (log2FC > 0.8 and adjusted *P* < 0.05) and downregulated genes (log2FC < −0.8 and adjusted *P* < 0.05) were analyzed separately using the enrichGO function. GO terms with an adjusted *P* value < 0.05 were considered statistically significant.

### Cell–cell communication analysis

Cell–cell communication analysis was performed using CellChat (v2.2.0)^54^ with the built-in mouse ligand–receptor database. Clusters containing fewer than 10 cells were excluded. Communication probabilities were inferred using the standard workflow based on the normalized gene expression matrix and cell-type annotations. The inferred MHC-I signaling from malignant cells to CD8^+^ T cells was then compared between treatment groups.

### TCR sequencing data analysis

T cell receptor (TCR) sequencing data were analyzed using contig-annotation files generated by DNBC4Tools. These files were integrated with cell barcode information from the scRNA-seq data to enable T-cell filtering, TCR clonotype assignment, clonotype quantification, and clonal expansion stratification.

First, cells in the contig-annotation files were matched to the scRNA-seq cell barcodes to link each TCR contig to its corresponding cell and cell-type annotation. Cells annotated as non-T cells were excluded, and only T cells were retained for downstream TCR analyses.

Within the filtered T-cell set, the clonotype supported by the highest read count in each cell was selected as the representative clonotype. Clonotypes were defined based on the TRB V(D)J gene segments (V, D, and J genes) and the complementarity-determining region 3 (CDR3) sequence. Clonotype frequency was calculated independently within each experimental group and defined as the proportion of cells carrying a given TRB clonotype.

Clonal expansion was quantified by stratifying TRB clonotypes into five categories according to clonotype frequency: Rare (≤ 0.01%), Small (0.01%–0.1%), Medium (0.1%–1%), Large (1%–10%), and Hyperexpanded (10%–100%)^55^. Shared TCR clonotypes were defined as clonotypes detected in both irradiated and distant tumors within the same treatment group. Shared expanded clonotypes were defined as shared clonotypes classified as Large or Hyperexpanded in at least one of the two tumor sites. All clonotypes were annotated using this grading scheme. For downstream differential analyses, a simplified two-tier classification was additionally generated: Large and Hyperexpanded clonotypes were grouped as expanded (Expanded), whereas Rare, Small, and Medium clonotypes were grouped as non-expanded (Non-expanded). Both the five-level and two-level classifications were retained for subsequent analyses.

### Pseudotime analysis

Pseudotime trajectories of CD8^+^ T cells and malignant cells were reconstructed using Monocle3 (v1.0.0)^56^. The normalized single-cell gene expression matrix was used for dimensionality reduction and trajectory graph learning, and an early-state population was manually selected as the root for each trajectory. For CD8^+^ T-cell trajectory analysis, CD8-Atypical, CD8-IFN/NK-like and Cycling populations were excluded. To functionally characterize CD8^+^ T-cell states along the trajectories, published T-cell functional gene signatures^57^ were scored in individual cells using the AddModuleScore function in Seurat. To compare the distributions of treatments along CD8^+^ T-cell pseudotime, cells were weighted to equalize the contribution of each pooled sample within each treatment group, and the weighted FA and CA distributions were estimated by Gaussian kernel smoothing; local FA–CA differences were expressed in percentage points. Pseudotime-associated genes along each CD8^+^ T-cell path were identified using the Monocle3 graph_test function, clustered according to their expression dynamics and subjected to functional enrichment analysis. Path-associated gene sets were scored using AUCell (v1.24.0) in cells assigned to the corresponding paths, and the resulting scores were visualized as box plots.

### Pathway activity scoring

All 50 Hallmark gene sets from the Molecular Signatures Database (MSigDB)^58^ were retrieved using the msigdbr R package (v7.5.1) and scored in individual malignant cells using UCell (v2.10.1)^59^. UCell scores were averaged within each malignant cell state and visualized as a heat map after row-wise Z-score normalization.

### Cellular modules analysis

The CoVarNet (v0.3.0)^29^ framework was applied to a cell-frequency matrix comprising annotated cell subtypes across samples. Non-negative matrix factorization identified 11 cellular modules (CMs), and module activities were calculated for each sample. Samples were assigned to a cellular module type (CMT) according to the CM with the highest activity. For each CM, CellChat-inferred interaction strengths among the top 15 contributing cell subsets were compared between samples assigned to the corresponding CMT and those assigned to all other CMTs.

### Flow cytometry

MC38 tumors were harvested on the days indicated in the figures and processed into single-cell suspensions using a tumor dissociation kit (Absin, Abs50090). Red blood cells were removed with RBC Lysis Buffer (BioLegend, 420301). For intracellular cytokine staining, cells were stimulated with a cell activation cocktail (BioLegend, 423303) for 4 h. Cells were then incubated with CD16/32 antibody (BioLegend, 101320) to prevent nonspecific binding, and live/dead discrimination was performed using the Zombie NIR™ Fixable Viability Kit (BioLegend, 423105). After that, cells were immunostained with fluorescent dye-conjugated antibodies against CD45 (1:100, BioLegend, 147704), CD8a (1:100, BioLegend, 100707), CD4 (1:100; BioLegend, 100412), F4/80 (1:100, BioLegend, 111604), IFN-γ (1:100, BioLegend, 505821), CD69 (1:100, BioLegend, 104505), γ-H2AX (1:50, BioLegend, 613404), Ly6G (1:100, BioLegend, 127613), CD11b (1:100, BioLegend, 101212), CD206 (1:100, BioLegend, 141708), and CD86 (1:100, BioLegend, 105014) on ice for 30 min. For intracellular staining, cells were fixed with Intracellular Staining Fixation Buffer (BioLegend, 420801) and then permeabilized with Intracellular Staining Permeabilization Wash Buffer (BioLegend, 421002) prior to antibody staining. Cells were acquired on a CytoFLEX flow cytometer (Beckman Coulter) and analyzed using FlowJo software (v10.8.1).

### Immunofluorescence

Neutrophils were first cultured in glass-bottom confocal dishes and treated as indicated in the figures. Before fixation, the cells were stained with SYTOX Green (1:10,000; Invitrogen, S7020) in Hanks’ Balanced Salt Solution (HBSS; Beyotime Inc., C0218) for 10 min at 37 °C to label extracellular DNA, and then fixed with 4% paraformaldehyde. For tumor tissues, samples were fixed in formalin, embedded in paraffin (FFPE), and sectioned into 5-μm slices by the histopathology platform. Antigen retrieval was performed using sodium citrate buffer (pH 6.0). Immunofluorescence staining was carried out following the Cell Signaling Technology protocol. Primary antibodies included anti-Cit H3 (1:2,000, Abcam, ab281584), anti-MPO (1:100, Abcam, ab90810), and directly conjugated AF488-anti-CD8a (1:100, Thermo Fisher Scientific, 53-0081-82) as indicated. After washing with PBST, secondary antibodies AF555 (1:1,000, Abcam, ab150114) and AF647 (1:1,000, Beyotime Inc., A0468) were added where needed. Nuclei were counterstained with DAPI (Abcam, ab104139). Finally, both slides and dishes were imaged on a Leica TCS SP8 confocal microscope.

### Multiplex immunofluorescence staining based on tyramide signal amplification (TSA)

The multiplex TSA fluorescence kit (ML0088-06, Nanjing Meladybio, China) was employed to label target proteins following the manufacturer’s instructions. The primary antibodies used were as follows: anti-CD8a (Abcam, ab217344), anti-CD4 (Abcam, ab183685), anti-granzyme B (Abcam, ab255598), anti-ISG15 (Abcam, ab315281), and anti-IFITM1 (Proteintech, 11727-3-AP). Image acquisition was performed using a multichannel fluorescence slide scanner (Pannoramic SCAN 150, 3DHISTECH). Due to the lower abundance of CD4⁺IFITM1⁺ cells relative to CD8⁺ISG15⁺ or CD8⁺granzyme B⁺ cells in the tumor microenvironment, we designated CD4⁺IFITM1⁺ cells as the reference population for spatial analysis. For each biological sample (n = 3 mice per group), 30 nearest-neighbor distance (NND) measurements were collected from 10 randomly selected fields of view. NND measurements were performed using the CaseViewer built-in measurement tool. To ensure statistical robustness and avoid pseudoreplication, the median NND of these 30 measurements was calculated per individual sample, and these median values were used for subsequent intergroup comparisons of the distances from CD4⁺IFITM1⁺ cells to CD8⁺ISG15⁺ or CD8⁺granzyme B⁺ cells, respectively.

### Immunohistochemistry for α-SMA

FFPE sections (5 μm) were deparaffinized, heated in sodium citrate buffer (pH 6.0) for antigen retrieval (15 min, pressure cooker), blocked with 3% H₂O₂ and 5% goat serum, then incubated with anti-α-SMA (1:100, Abcam, ab7817) overnight at 4 °C. After HRP-secondary antibody (1 h, room temperature), DAB was used for signal development, followed by hematoxylin counterstaining. Images were captured with a digital slide scanner (Grundium, Ocus). The percentage of α-SMA-positive area was quantified in 5 random fields per sample using the IHC Profiler plugin in ImageJ (v1.53c), which automatically classifies stained pixels into high positive, positive, low positive, and negative categories based on optical density.

### Mouse neutrophils isolation and treatment

Mouse neutrophils were immunomagnetically isolated from the bone marrow of C57BL/6J mice (6–8 weeks old) using the Ly6G microbead kit (Miltenyi, 130-120-337) according to the manufacturer’s instructions. To assess NET formation induced by tumor-derived factors, neutrophils (2 × 10⁵) were resuspended in HBSS and seeded into glass-bottom confocal dishes. Conditioned media were collected from MC38 cells at 24 h after no irradiation, 8-Gy CONV irradiation, or 8-Gy FLASH irradiation, and then added to the neutrophils for co-culture. After 4 h of incubation at 37 °C and 5% CO₂, NET formation was evaluated by immunofluorescence. Separately, for inhibition studies, GSK484 (10 μmol/L) or PJ34 (20 μmol/L) was added to cultured neutrophils 30 min before irradiation (8-Gy CONV or FLASH). After irradiation, cells were incubated at 37 °C and 5% CO₂ for 4 h, and then NET formation was assessed by immunofluorescence. PMA (1 nmol/L, TargetMol, TQ0198) was added to induce NETosis as a positive control.

### ROS detection

Neutrophils were incubated with the ROS indicator CM-H₂DCFDA (1:1,000, Beyotime Inc., S0036S) diluted in HBSS. Immediately after probe addition, cells were irradiated with 8-Gy X-rays delivered at either CONV or FLASH dose rate. Following irradiation, cells were incubated at 37 °C for 30 min, and then the fluorescence intensity in the FITC channel was measured by flow cytometry. As a positive control, neutrophils were treated with 2 nM PMA (instead of irradiation) to induce ROS production. All steps were performed in the dark to minimize photobleaching.

### Immunoblotting

Cells were lysed in RIPA buffer (Beyotime Inc., P0013B) with protease inhibitor cocktail (Bimake, B14001) on ice. Proteins were separated by SDS-PAGE, transferred to PVDF membranes (Millipore, ISEQ00010), blocked, and probed with primary antibodies followed by HRP-conjugated secondary antibodies. Signal was detected using the Tanon 4600 chemiluminescence imaging system (Tanon Science & Technology Co., Ltd., Shanghai, China). The primary antibodies used were as follows: anti-Histone H3 (1:1,000, CST, 4499), anti-Acetyl-Histone H3 (1:2,000, Active Motif, 39140), anti-H3K9ac (1:2,000, Active Motif, 39918), anti-H3K27ac (1:2,000, Active Motif, 39134), anti-H3K14ac (1:2,000, Active Motif, 65702), anti-H3K56ac (1:2,000, Active Motif, 61061), anti-H4K5ac (1:2,000, Active Motif, 39700), anti-poly/mono-ADP ribose (1:1,000, CST, 89190), and anti-GAPDH (1:1,000, Beyotime Inc., AF0006). Protein levels were quantified using the ImageJ software (v1.53c).

### Mouse bone marrow derived-macrophages (BMDMs) culture, polarization, and irradiation

Bone marrow cells were flushed from mouse femurs and tibiae, treated with red blood cell lysis buffer, and then resuspended in DMEM complete medium for 4 h. Non-adherent cells were collected, resuspended in RPMI 1640 medium supplemented with 10% FBS, 2% penicillin-streptomycin, and 25 ng/mL M-CSF (Novoprotein, CB34), and seeded into 6-well plates. The day of seeding was designated day 1. Fresh M-CSF-containing medium was added on day 3 and replaced on day 5. On day 6, BMDMs were polarized to M1 with IFN-γ (50 ng/mL, Novoprotein, C746) plus LPS (50 ng/mL, Selleck, S7850), or to M2 with IL-4 (10 ng/mL, Novoprotein, CK15) plus IL-13 (10 ng/mL, Peprotech, 210-13), for 24–48 h. At 24 h after cytokine addition, cells were irradiated with 8-Gy CONV X-rays. Samples were collected before irradiation and at 3, 6, 12, and 24 h post-irradiation for immunoblotting. For flow cytometry analysis of polarization, cells were harvested 24 h after cytokine stimulation.

### Statistical analysis

Data are presented as mean ± SEM. Normality was assessed by the Shapiro–Wilk test, and variance homogeneity was assessed by the F test (for two groups) or Levene”s test (for multiple groups). For two-group comparisons, an unpaired two-tailed Student’s *t*-test was used when variances were equal; otherwise, Welch’s *t*-test was applied. For multiple groups, one-way ANOVA followed by Tukey’s HSD test was performed. Tumor growth was analyzed by repeated-measures ANOVA with treatment, time, and their interaction as factors, followed by Tukey’s post hoc test for pairwise comparisons. Sphericity was assessed with Mauchly’s test and corrected using the Greenhouse–Geisser correction where applicable. Survival curves were estimated by the Kaplan–Meier method and compared by the log-rank test. A *P* value < 0.05 was considered significant. Sample sizes were determined on the basis of preliminary experiments and prior experience, and are indicated in the figure legends. No blinding was used in this study. All quantitative analyses were performed according to predefined, objective criteria to minimize observer bias, and automated quantification was used where possible. Mice were randomly assigned to treatment groups using a random number table. To ensure the reproducibility of our results, all experiments were conducted with at least twice independent replicates. All statistical analyses were performed using GraphPad Prism 9.3.0.

## Data availability

The sequencing data supporting our findings have been deposited into the China National GeneBank Sequence Archive (CNSA) of CNGBdb with accession number CNP0009637 and Cancer Omics Atlas of Spatiotemporal Topology (COAST) database with accession number of COA0000020. All data supporting the findings of this study are available within the Article and its Supplementary Information. Source data are provided with this paper.

## Acknowledgments

We acknowledge Drs. Z. Chai, Tom. K. Hei, S. Wang, F. Liu, J. Diwu, H. Shi, J. Yu from the School of Radiation Medicine and Protection, Soochow University, for their support. We acknowledge Zhongjiu Flash Medical Technology Co., Ltd. for providing the FLASH irradiation device, and we thank H. Feng, Y. Jiang, Y. Luo, Q. Yang, L. Zhou, P. Wu, K. Gan, D. Cheng, and W. Su for provided technical support for the operation of the FLASH irradiation device during the FLASH irradiation experiments. We acknowledge P. Zhang, L. Chen, C. Lao, J. Liu, T. He, and S. Wang from the Institute of Applied Electronics, China Academy of Engineering Physics, for their support during FLASH X-rays experiments. We acknowledge B. Lin and X. Mao from Department of Oncology, Nuclear Medicine Laboratory of Mianyang Central Hospital, for their support during CONV X-rays experiments. We acknowledge Y. Xu, from the Department of Biochemistry, School of Integrative Medicine, Shanghai University of Traditional Chinese Medicine, and C. Di, from the Institute of Modern Physics, Chinese Academy of Sciences, for their valuable discussions. We acknowledge State Key Laboratory of Genome and Multi-omics Technologies, BGI Research, for their support, and we thank Y. Zhang for provided technical support in the single cell experiment. We acknowledge the Collaborative Innovation Center of Radiological Medicine of Jiangsu Higher Education Institutions, the Jiangsu Key Laboratory of Radiation Injury Prevention and Hazardous Substance Control, the Key Laboratory of Radiation Damage and Treatment of Jiangsu Provincial Universities and Colleges, and the project funded by the Priority Academic Program Development of Jiangsu Higher Education Institutions (PAPD) for their support.

## Author contributions

Conceptualization: H. Z., C. S., D. W., M. X., Y. Y., Jianxin W.; Data curation: X. S., X. Ding., Q. L., H. L.; Formal analysis: X. S., S. Q., Y. J., M. L. and B. L.; Funding acquisition: H. Z., C. S., D. W., Y. Y., Jianxin W., X. S.; Investigation: X. S., X. Ding, S. Q., C. C., Y. H., G. S., F. J., Y. X., H. Q., X. J., T. W.; Methodology: X. S., Y. Y., Jianxin W., Q. S.; Project administration: C. Ye, C. Yan, J. Zhang, J. H., Y. P., Y. B., Z. Z., K. Z., D. X., P. L., C. C., L. S., Y. L., F. G., L. W.; Resources: H. Z., S. C., W. D., M. X., X. Du, J. Zhao, F. X., Jian W.; Supervision: H. Z., S. C., W. D., M. X.; Validation: X. S., S. Q., Y. Y., Y. J., Jianxin W.; Visualization: X. S., S. Q., Y. Y., Y. J., Y. H., Y. W.; Writing – original draft: H. Z., C.S., X. S., S. Q., Y. J.; Writing – review & editing: All authors contributed to the final version of the paper.

---

**Extended Data Fig. 1.**
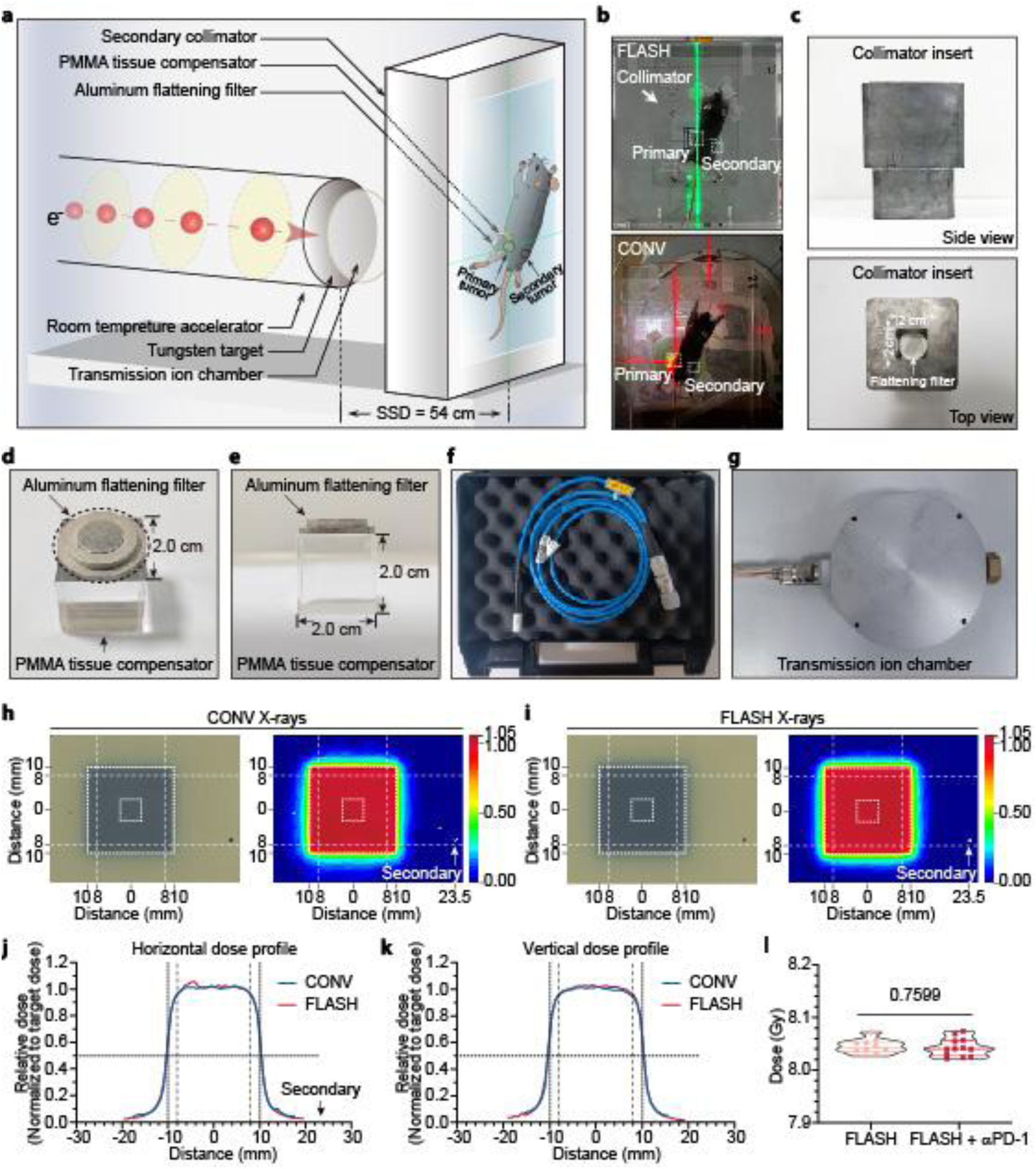
Irradiation device, setup and dosimetric analysis. **a**, Schematic diagram of X-ray FLASH irradiation. **b**, The positioning of the tumor bearing mice exposed to FLASH or CONV X-rays is shown. An 8-Gy horizontal beam was delivered to the primary tumor on day 10 post-implantation. **c**, Lead secondary collimators with an aperture size of 2 × 2 cm were used to delimit the FLASH irradiation field for the MC38 tumor. **d**,**e**, An aluminum flattening filter was attached to a 2 × 2 × 2 cm PMMA tissue compensator, combining dose buildup and lateral profile homogenization. **f**, A PTW diamond detector was used to measure the dose at the field center (5 × 5 mm). **g**, A custom-built low-pressure ionization chamber (transmission ionization chamber) was used for real-time monitoring of dose and spatial distribution during FLASH irradiation. **h**,**i**, EBT4 films were used for dose evaluation; the corresponding dose distributions from CONV (**h**) and FLASH (**i**) irradiation are shown. The location of the secondary tumor is indicated by arrows. **j**,**k**, Horizontal (**j**) and vertical (**k**) dose profiles extracted from the distributions shown in (**h**) and (**i**), respectively. **l**, Individual absorbed doses in mice receiving FLASH X-rays alone or with αPD-1; statistical significance was determined by unpaired two-tailed Student’s *t*-test.

**Extended Data Fig. 2.**
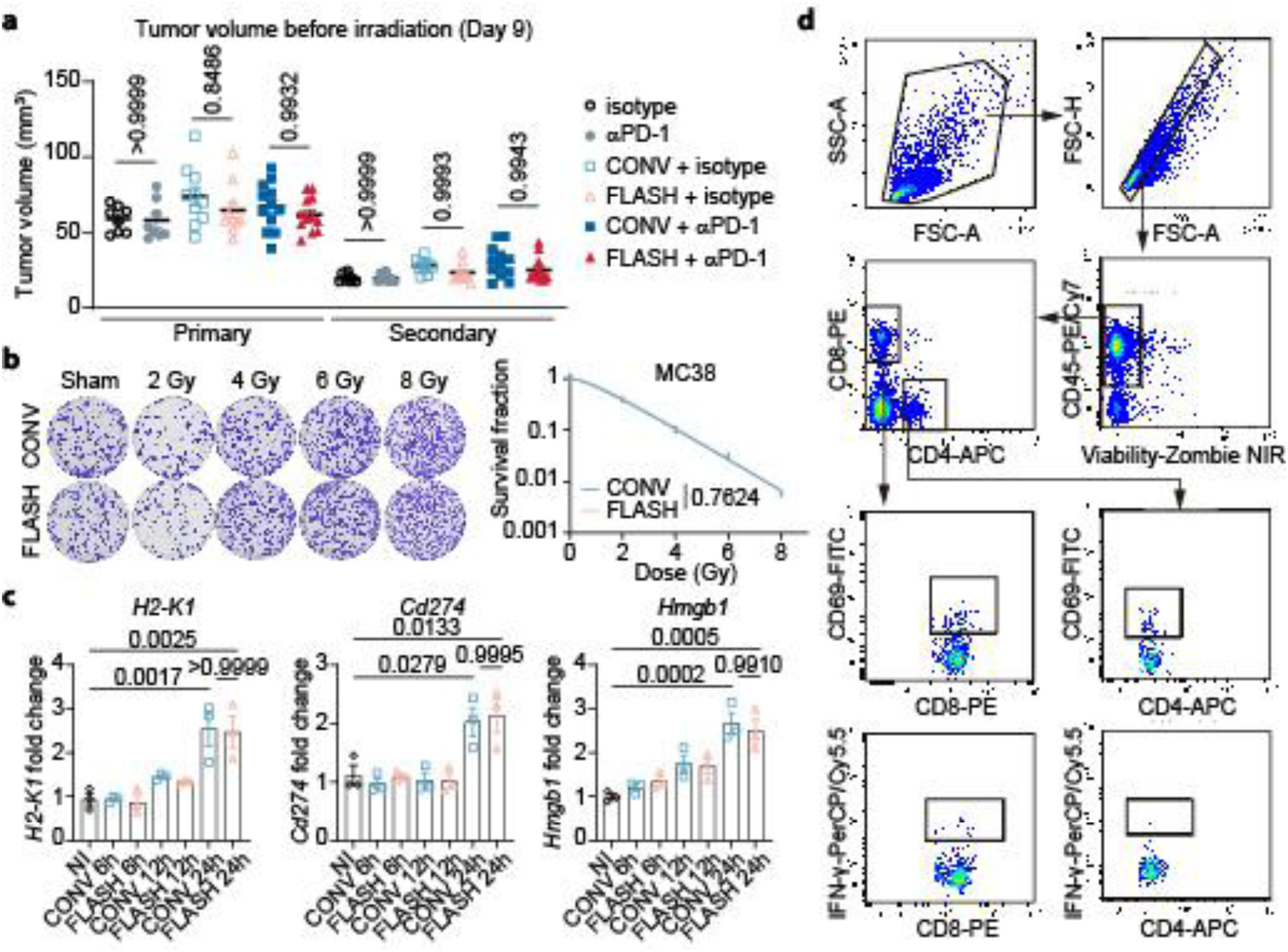
Characterization of baseline tumor burdens and *in vitro* responses to FLASH and CONV irradiation. **a**, Tumor volume of the indicated groups on the day before irradiation. **b**, Representative images of colonies formed in sham-irradiated, CONV-irradiated, and FLASH-irradiated groups at the indicated doses. Survival curves were fitted using the single-hit multi-target model, and data points represent the mean ± SEM from three independent experiments. **c**, Relative mRNA expression of *H2-K1*, *CD274*, and *Hmgb1* in MC38 cells after 8-Gy CONV or FLASH irradiation. Expression was measured by quantitative real-time PCR (qRT-PCR) and normalized to the non-irradiated (NI) group. Data are presented as mean ± SEM (n = 3 per group). **d**, Gating strategies for identifying CD69^+^ and IFN-γ^+^ populations in CD4^+^ and CD8^+^ T cells from the splenocytes described in Fig. 1j. Statistical significance was determined by one-way ANOVA (**a**,**c**) and two-way ANOVA (**b**).

**Extended Data Fig. 3.**
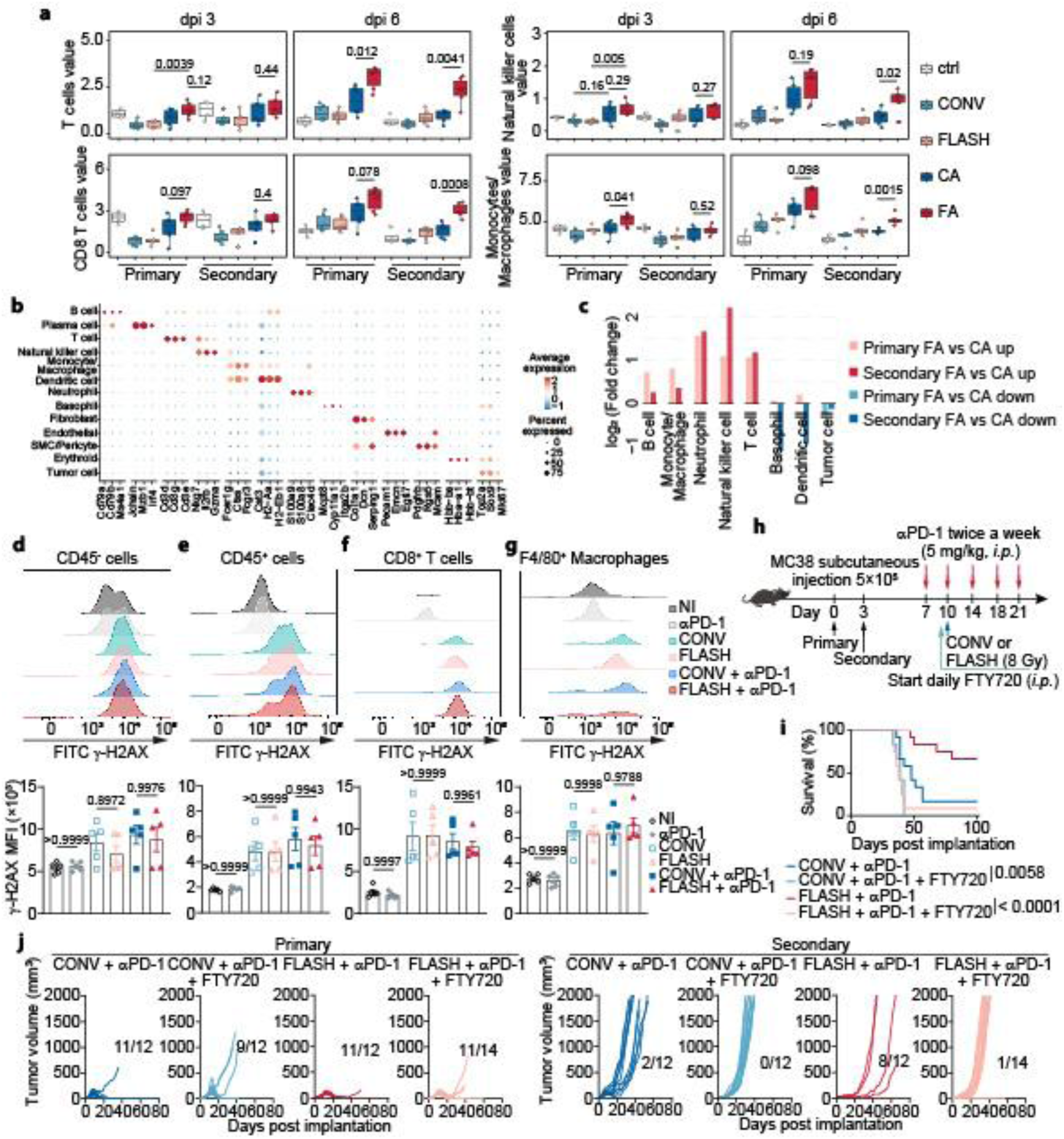
Assessment of intratumoral immune infiltration, DNA damage, and T-cell migration following FLASH and CONV X-rays. **a**, Box plots showing the relative abundance of T cells, CD8^+^ T cells, natural killer cells, and monocytes/macrophages estimated by mMCP-counter across the different treatment groups. **b**, Dot plot showing the expression of signature genes across the major cell types. Color denotes scaled average expression and dot size denotes the fraction of cells expressing each gene. **c**, Fold change in the abundance of major cell lineages in irradiated and distant tumors in the FLASH X-rays plus αPD-1 (FA) versus CONV X-rays plus αPD-1 (CA) groups. **d**–**g**, Evaluation of DNA double-strand breaks (DSBs) in the primary tumor. Tumor-bearing mice were assigned to the indicated groups. At 6 h post-irradiation, tumors were harvested and dissociated into single-cell suspensions, and γ-H2AX was measured in CD45^−^ (**d**), CD45⁺ (**e**), CD8⁺ (**f**), and F4/80⁺ (**g**) cells. Data are shown as mean ± SEM (n = 5 mice per group). **h**, Schema of the experimental strategy to evaluate the impact of inhibiting T-cell egress from lymphoid tissues on the anti-tumor effect induced by FLASH X-ray combined with αPD-1. **i**,**j**, Tumor growth and mouse survival are evaluated in the FA and CA groups treated with or without FTY720. Primary and secondary tumor volumes of individual mice are plotted, and the ratio of tumor-free mice to total mice is indicated for each treatment group (**i**). Kaplan–Meier survival curves are shown in (**j**). Statistical significance was assessed using unpaired two-tailed Student’s *t*-test (**a**), one-way ANOVA (**d**–**g**), and log-rank test (**j**). NI, Non-irradiated.

**Extended Data Fig. 4.**
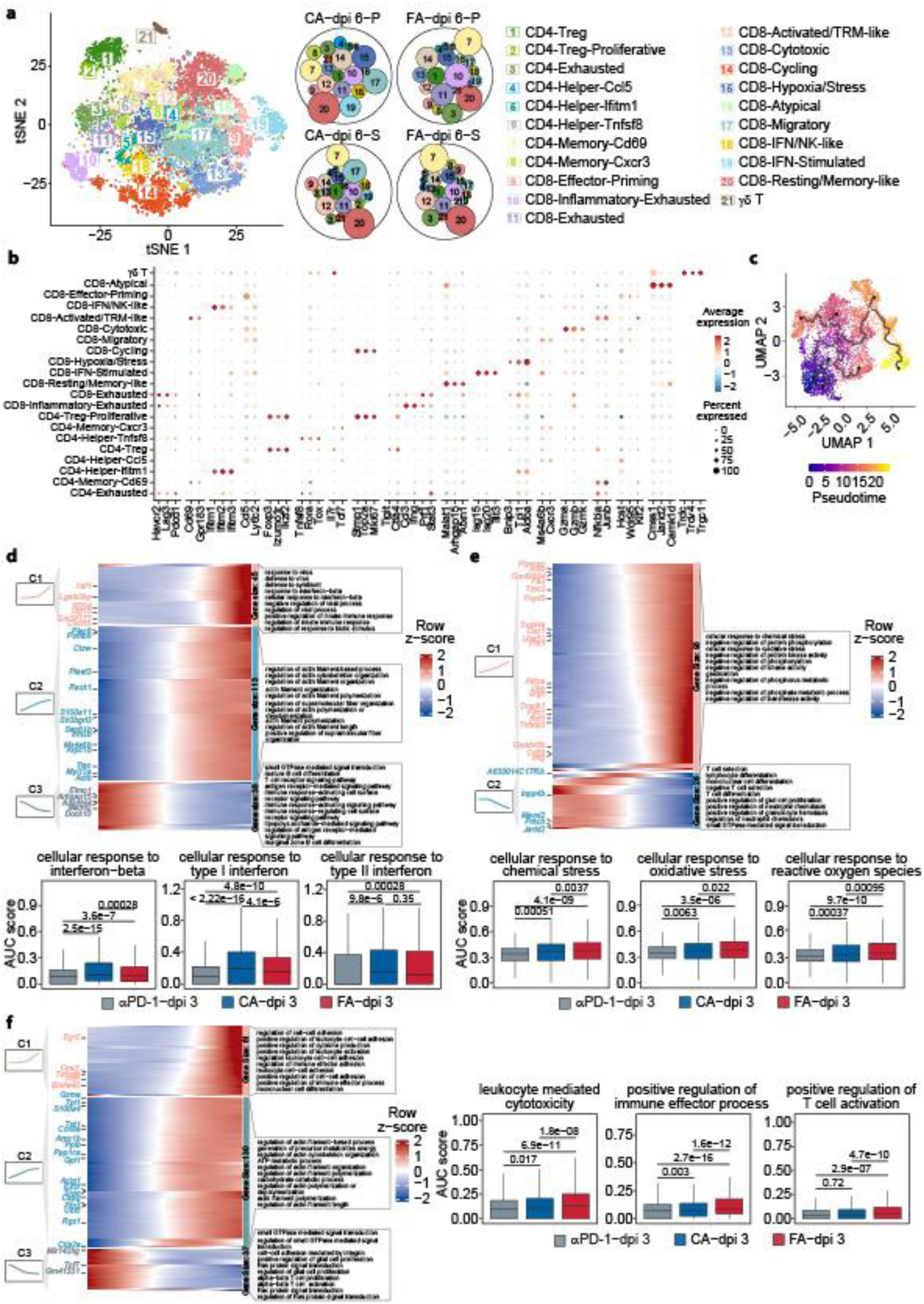
Subclustering and pseudotime trajectory analysis of CD8^+^ T subsets. **a**, UMAP of T cells from αPD-1, FLASH + αPD-1, and CONV + αPD-1 groups, colored by 21 T-cell subclusters; a bubble plot shows the proportions of the 21 T-cell subclusters across the indicated conditions. **b**, Dot plot showing the expression of signature genes across the 21 T-cell subclusters. Color denotes scaled average expression and dot size denotes the fraction of cells expressing each gene. **c**, Diffusion map of CD8^+^ T cells colored by pseudotime. **d**–**f**, Heat map showing scaled expression of genes differentially expressed along pseudotime in CD8^+^ T cells assigned to path 1 (**d**), path 2 (**e**) and path 3 (**f**). Enriched pathways associated with pseudotime-dependent gene modules are annotated alongside the heat maps, and adjacent box plots show AUC scores of selected pathways across the αPD-1, CA and FA groups. Significance was determined by Wilcoxon test (**d**–**f**).

**Extended Data Fig. 5.**
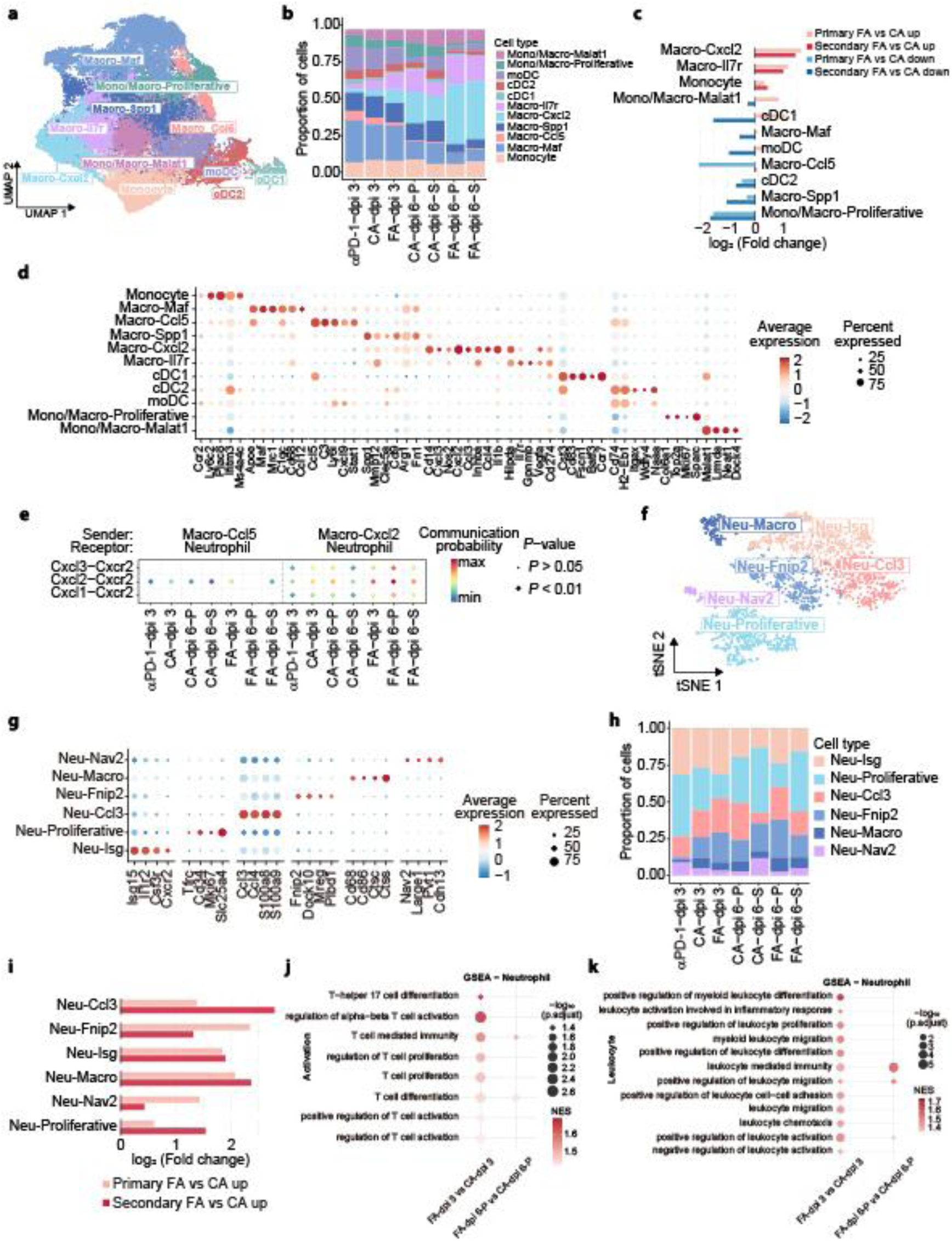
Comparison of intratumoral myeloid subsets between FA and CA tumors by single-cell analysis. **a**, UMAP of mononuclear phagocyte single cells from tumors collected on days 3 and 6 post-irradiation, colored by 11 subsets. **b**, Proportions of the 11 mononuclear phagocyte subsets across the indicated groups. **c**, Fold change in the abundance of the 11 mononuclear phagocyte subsets in irradiated and distant tumors in the FA versus CA groups. **d**, Dot plot showing the expression of signature genes across the 11 mononuclear phagocyte subsets. **e**, Bubble plot showing significant CXCL-CXCR2 chemokine interactions between the selected macrophage subsets and neutrophils. **f**, UMAP of neutrophils from tumors collected on days 3 and 6 post-irradiation, colored by six subsets. **g**, Dot plot showing the expression of signature genes across the six neutrophil subsets. **h**, Proportions of the six neutrophil subsets across the indicated groups. **i**, Fold change in the abundance of the six neutrophil subsets in irradiated and distant tumors in the FA versus CA groups. **j**, GSEA bubble plot comparing T-cell-related pathways in neutrophils between FA and CA across post-irradiation time points and tumor sites. **k**, GSEA bubble plot comparing myeloid/leukocyte-associated pathways in neutrophils between FA and CA across post-irradiation time points and tumor sites. For **d**,**e**, color denotes scaled average expression and dot size denotes the fraction of cells expressing each gene. For **j**,**k**, positive NES values indicate enrichment in the FA versus CA groups; point size denotes –log_10_ (adjusted *P* value). dpi, days post irradiation.

**Extended Data Fig. 6.**
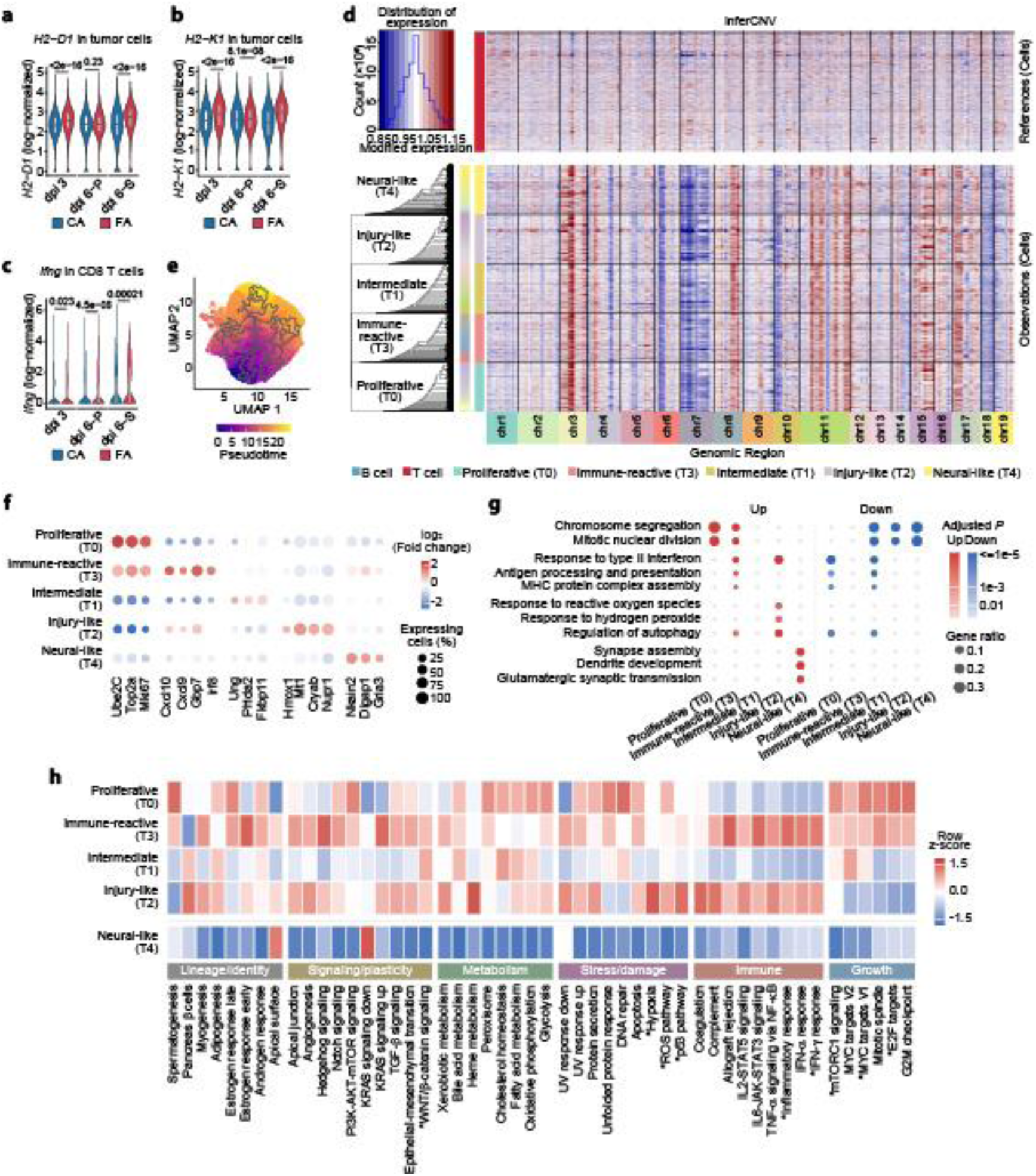
Single-cell landscape of malignant-cell heterogeneity and MHC-I expression. **a**–**c**, Violin plots showing the expression of *H2-D1* (**a**), *H2-K1* (**b**) in tumor cells, and *Ifng* (**c**) in CD8^+^ T cells from scRNA-seq data. **d**, inferCNV analysis of malignant cell states. Copy-number variation (CNV) profiles of the five malignant cell states (T0–T4) were inferred using inferCNV with B and T cells as normal references cells. Red and blue denote putative copy-number gains and losses, respectively, across chromosomes 1–19. **e**, Diffusion map of malignant cells colored by pseudotime. **f**, Dot plot of representative marker genes across the five malignant cell states (T0–T4). Color denotes average log_2_ fold change and dot size denotes the fraction of cells expressing each gene. **g**, Gene Ontology enrichment of upregulated and downregulated genes in each malignant cell state (T0–T4). Dot size denotes gene ratio and color denotes adjusted *P* value. **h**, Heat map of mean UCell scores for Hallmark gene sets across the five malignant cell states (T0–T4). Values are row Z-scores. Asterisks indicate gene sets selected for presentation in Fig. 3d. Significance was determined by Wilcoxon test (**a**–**c**).

**Extended Data Fig. 7.**
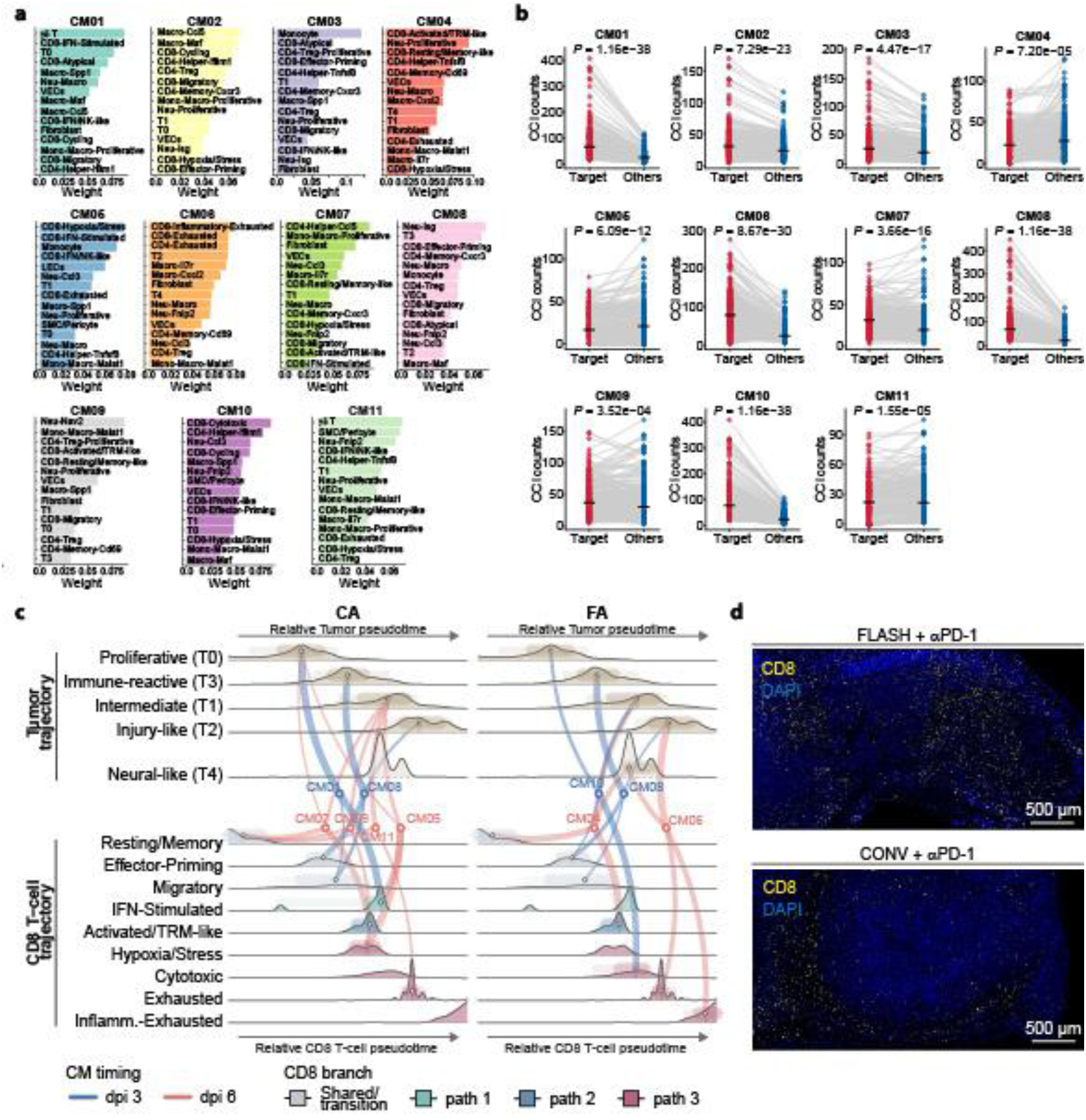
Cellular module composition, cell–cell interaction strength, and tumor–CD8^+^ T-cell coupling under FA versus CA. **a**, Cellular composition of the 11 CoVarNet CMs. For each CM, the 15 cell subsets with the highest non-negative matrix factorization (NMF) weights are shown; bar length denotes NMF weight. **b**, Comparison of cell–cell interaction (CCI) strengths for CM-associated cell subsets between samples assigned to the corresponding CMT and the remaining samples. Each dot represents one cell-subset pair within a CM; paired values for the same cell subset pair are connected by lines. **c**, Coupling of malignant cell states and CD8^+^ T-cell states within treatment-associated CMs under CA and FA. For each CM, the two highest-weight malignant cell states and the two highest-weight CD8⁺ T-cell states are positioned according to their relative pseudotime along the corresponding malignant cell and CD8^+^ T-cell trajectories and connected by lines. Line color denotes CM timing (day 3 or day 6 post-irradiation), line thickness denotes NMF weight, and CD8^+^ T-cell state color denotes trajectory branch (paths 1–3). **d**, Representative images show the spatial distribution of CD8⁺ T cells (yellow) within irradiated tumors, corresponding to Fig. 3i (scale bar, 500 μm). Significance was determined by Wilcoxon test (**b**).

**Extended Data Fig. 8.**
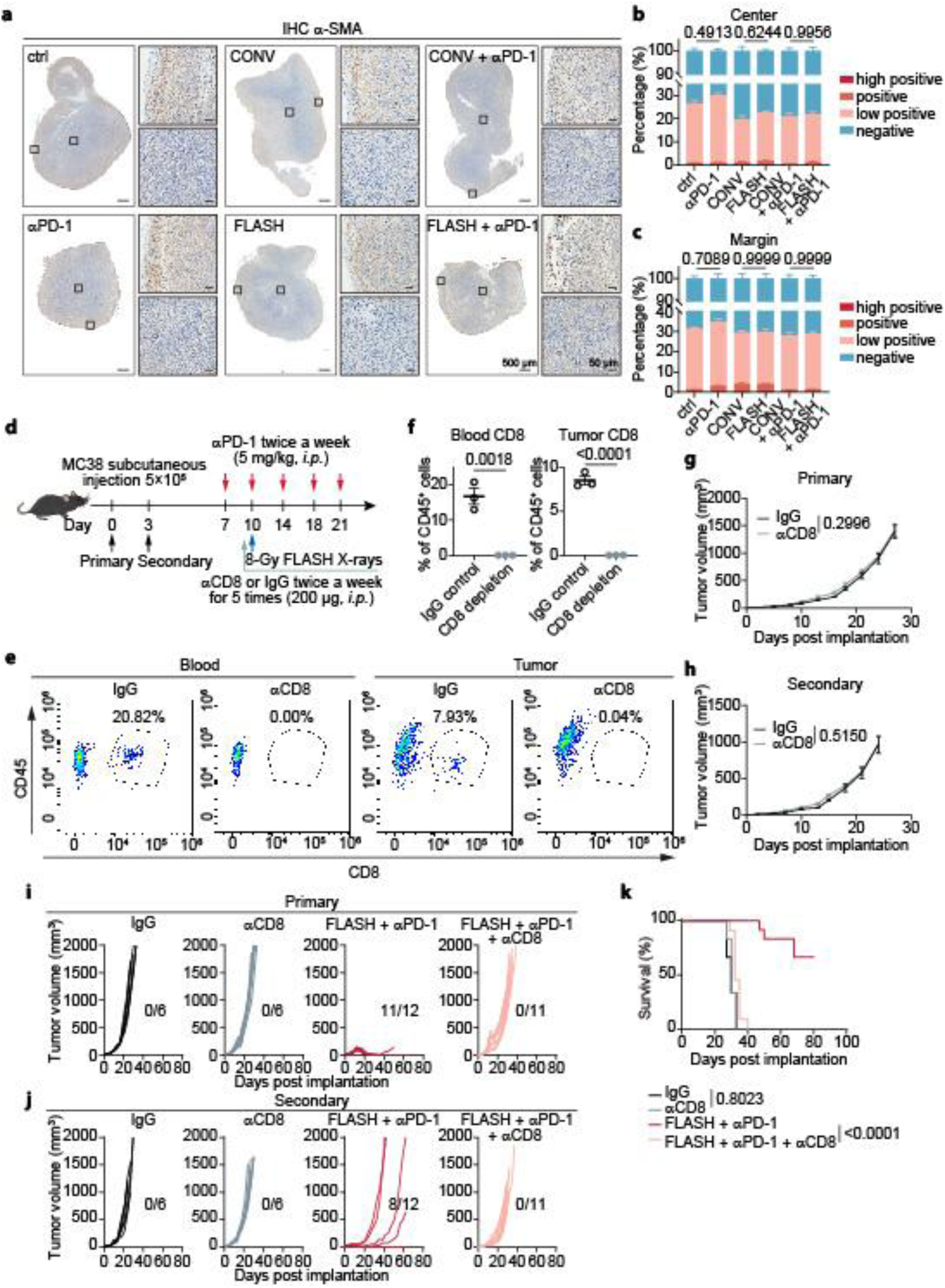
Alpha-smooth muscle actin (α-SMA) expression and antitumor responses following CD8^+^ T-cell depletion. **a**–**c**, α-SMA expression was assessed by immunohistochemistry of primary tumors from the indicated groups at day 3 post-irradiation (**a**). The percentages of high-positive, positive, low-positive, and negative α-SMA staining were quantified using the IHC Profiler plugin in ImageJ and are shown for the tumor center (**b**) and tumor margin (**c**) (n = 6 mice per group). **d**, Schema for evaluating the role of CD8⁺ T cells in the abscopal effect induced by FLASH X-rays combined with αPD-1. **e**–**h**, Evaluation of CD8⁺ T-cell depletion efficiency by the αCD8 antibody and its effect on tumor growth. Representative flow cytometry plots (**e**) and quantification (**f**) of CD8⁺ T-cell abundance in peripheral blood and tumors from IgG-versus αCD8-treated mice are shown. Primary (**g**) and secondary (**h**) tumor growth curves are plotted for both groups. **i**–**k**, Mouse survival and tumor growth are evaluated in the indicated groups. Primary (**i**) and secondary (**j**) tumor volumes of individual mice are plotted, and the ratios (tumor-free mice/total mice) are indicated for each treatment group. Kaplan–Meier survival curves are shown in (**k**). Data are presented as mean ± SEM. Statistical significance was assessed using one-way ANOVA (**b**,**c**), unpaired two-tailed Student’s *t*-test (**f**), repeated measures ANOVA (**g**,**h**), and log-rank test (**k**).

**Extended Data Fig. 9.**
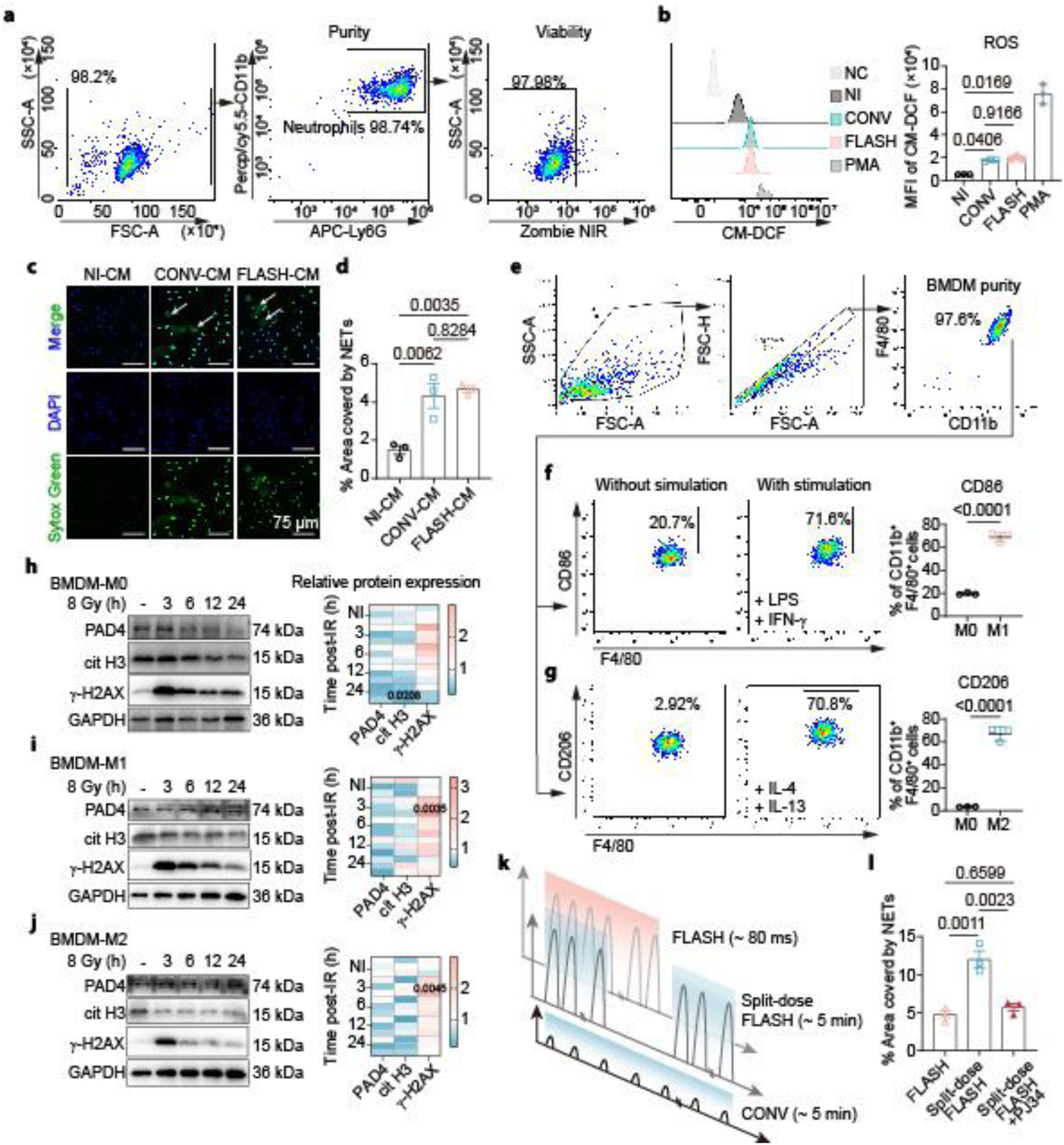
*In vitro* responses of neutrophils and macrophages to CONV and FLASH X-rays. **a**, Neutrophils were immunomagnetically isolated from the bone marrow of mice using the Ly6G microbead. The viability and purity were assessed by flow cytometry. **b**, Intracellular ROS levels in neutrophils were detected at 30 min post-irradiation using the ROS probe CM-H₂DCFDA. The fluorescence intensity of CM-DCF (reflecting ROS levels) was measured by flow cytometry and quantified. **c**,**d**, Neutrophils were co-cultured with conditioned media (CM) collected from MC38 cells subjected to 8-Gy irradiation. After 4 h, NET release was visualized by SYTOX Green labeling of extracellular DNA (**c**) and quantified as the percentage of NET-covered area (**d**). **e**, Schematic representation of the split-dose FLASH irradiation regimen. A single FLASH macro-pulse was equally split into two macro-pulses, sequentially delivering a total X-ray dose of 8 Gy (4 Gy per pulse). **f**, Quantification of NETs corresponding to the data presented in Fig. 5k. **g**–**l**, Bone marrow cells were treated with M-CSF to generate BMDMs (**g**), and then polarized into M1 with IFN-γ plus LPS (**h**) or into M2 with IL-4 plus IL-13 (**i**). The expression levels of PAD4, Cit H3, and γ-H2AX at 3, 6, 12, and 24 h post-irradiation were determined by immunoblotting and quantified in M0 (**j**), M1 (**k**), and M2 (**l**) cells. Statistical significance was assessed using one-way ANOVA (**b**,**d**,**f**,**j**–**l**), and the unpaired two-tailed Student’s *t*-test (**h**,**i**). NI, Non-irradiated.

**Extended Data Fig. 10.**
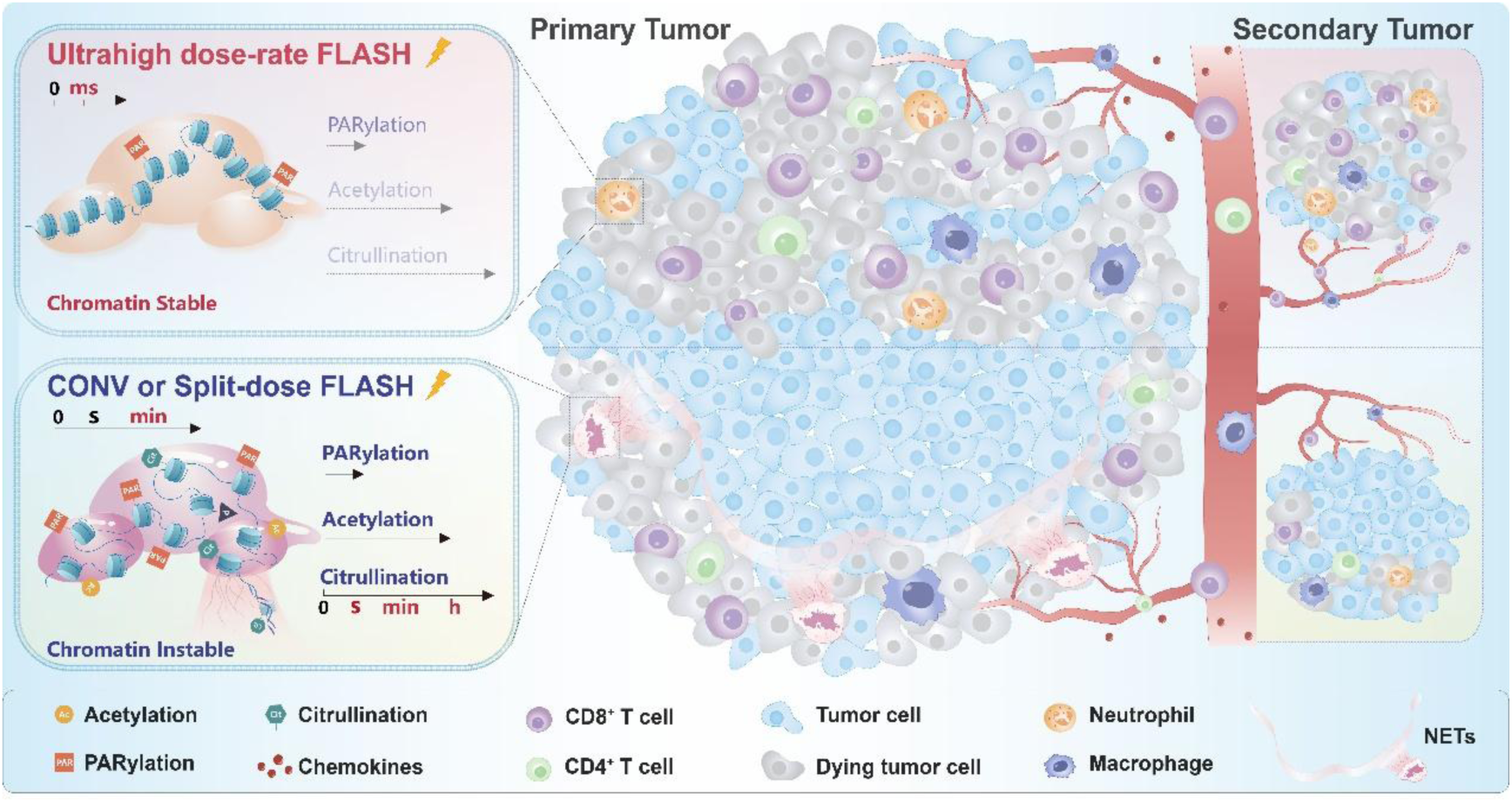
Proposed mechanism of FLASH-induced abscopal effect through neutrophil chromatin preservation. FLASH X-rays delivered within milliseconds prevent time-dependent chromatin decondensation and histone acetylation in neutrophils, thereby suppressing NETosis and dismantling the physical barriers built by NETs. This allows deep infiltration of CD8⁺ T cells within the irradiated tumor and promotes T-cell clonal expansion as well as T-cell receptor (TCR) sharing across tumor sites, thereby conferring a potent abscopal effect. In contrast, CONV X-rays delivered over minutes permit full execution of chromatin remodeling and histone acetylation, leading to massive NET formation. These NETs physically block CD8⁺ T cells from entering the tumor center and restrict their clonal expansion, ultimately resulting in the failure of CONV combined with αPD-1 to induce a robust abscopal effect.

